# *Atrx* deletion impairs cGAS-STING signaling and increases response to radiation and oncolytic herpesvirus in sarcoma

**DOI:** 10.1101/2021.03.08.434225

**Authors:** Warren Floyd, Matthew Pierpoint, Chang Su, Lixia Luo, Katherine Deland, Amy J. Wisdom, Daniel Zhu, Yan Ma, Suzanne Bartholf DeWitt, Nerissa T. Williams, Jason A. Somarelli, David L. Corcoran, William C. Eward, Diana M. Cardona, David G. Kirsch

## Abstract

*ATRX* is one of the most frequently altered genes in solid tumors, and mutation is especially frequent in soft tissue sarcomas. However, the role of *ATRX* in tumor development and response to cancer therapies remains poorly understood. Here, we developed a primary mouse model of soft tissue sarcoma and showed that *Atrx* deleted tumors are more sensitive to radiation therapy. In the absence of *Atrx*, irradiated sarcomas have increased persistent DNA damage, telomere dysfunction, and mitotic catastrophe. We find that *Atrx* deleted sarcomas have a reduced adaptive immune response, impaired cGAS-STING signaling, and increased sensitivity to an oncolytic herpesvirus therapy. Translation of these results to patients with *ATRX* mutant cancers could enable genomically-guided cancer therapeutic approaches that improve patient outcomes.

## Introduction

*ATRX* (alpha-thalassemia/mental retardation, X-linked) is a chromatin remodeling protein and tumor suppressor, with mutations or copy number alterations occurring in approximately 6% of the more than 65,163 tumors sequenced in the AACR GENIE database (1). Mutations in *ATRX* are predominantly loss of function, and are enriched in specific cancer types, including low grade gliomas, soft tissue sarcomas, pancreatic neuroendocrine tumors, and uterine corpus endometrial carcinomas (2–5). The effect of *ATRX* on overall survival (OS) varies by cancer type. *ATRX* mutation is associated with improved OS in gliomas, but worse OS in pancreatic neuroendocrine tumors (6, 7).

ATRX is hypothesized to play key roles in both normal and cancer cells, such as global epigenetic maintenance of pericentromeric heterochromatin, repression of alternative lengthening of telomeres (ALT), targeting of polycomb repressor complex 2, and regulation of DNA double strand break (DSB) damage repair (8–10). In the context of cancer therapies, previous work suggests that *Atrx* deletion impairs the DNA damage response, though no single mechanism has been identified (11–13). There are also conflicting reports regarding the impact of *Atrx* deletion on radiation response, with increased radiosensitivity reported in some, but not all, studies (10, 11). Further work has demonstrated that *Atrx* plays an important role in maintaining telomere genomic stability in embryonic stem cells and in neuroprogenitor cells (10).

Interestingly, recent reports have identified the ATRX as an important modulator of the cyclic guanosine monophosphate-adenosine monophosphate synthase (cGAS) and its adapter protein Stimulator of IFN Gene (STING) pathway’s response to extrachromosomal telomeric repeat DNA in cancer cell lines with alternative lengthening of telomeres (ALT) (14). The cGAS-STING pathway acts as an innate immune sensor for microbial and viral pathogens by detecting double stranded DNA (dsDNA) in the cytoplasm of mammalian cells, and has emerged as an important link between DNA damage and the innate immune system (15–17). Engagement of the cGAS-STING pathway results in the phosphorylation of IFN regulatory factor 3 (IRF3) and transcriptional induction of type-I interferons (18).

To study the role of *Atrx* in cancer development and therapeutic response, we generated a primary genetically engineered mouse model of soft tissue sarcoma with and without *Atrx* deletion. We found that *Atrx* deletion increases radiation-induced persistent DNA damage, mitotic dysfunction, and radiotherapy response in both immune proficient and T-cell deficient mouse models. We further show that *Atrx* deleted primary sarcomas have impaired adaptive immune response and reduced tumor-intrinsic cGAS-STING signaling after radiation. Finally, we demonstrate that sarcomas with *Atrx* deletion and aberrant cGAS-STING signaling are sensitized to oncolytic herpesvirus therapy. Taken together, these findings have important implications for precision oncology in *ATRX*-mutant cancers.

## Results

To model *ATRX* alterations from human cancer in a primary mouse model, we examined The Cancer Genome Atlas (TCGA), a large next generation sequencing cancer database(4). Based on the TCGA database we determined that three of the most common subtypes of soft tissue sarcoma, undifferentiated pleomorphic sarcoma, myofibrosarcoma, and leiomyosarcoma (STS cohort), had recurrent alterations in the *ATRX* gene (24% of samples). The majority of these ATRX alterations were copy number deletions, frameshift mutations, or missense mutations located within the gene’s functional domains (Figure 1A-D) (5). We therefore concluded that *ATRX* alterations in human cancer can faithfully be modeled via conditional deletion, and that soft tissue sarcoma is a relevant model system in which to study the impact of ATRX loss-of-function in cancer. Next, we found that within this STS cohort in the TCGA database, ATRX alteration was associated with significantly worse disease specific survival (Supplemental Figure 1A). This ATRX associated worse disease specific survival was even more pronounced in tumors that did not receive ionizing radiation (Figure 1E). Interestingly however, survival was similar for patients with sarcomas with and without ATRX alterations if they were treated with radiation therapy (Figure 1F), suggesting that ATRX loss of function may increase the radiation response of soft tissue sarcomas. Furthermore, we examined a publicly available data set of whole genome sequencing from human undifferentiated pleomorphic sarcoma (UPS), one of the most commonly diagnosed subtypes of soft tissue sarcoma in adults (2). Analysis of this dataset revealed that human UPS with *ATRX* alterations, had a significant increase in chromosomal rearrangements and an increased likelihood of SBS3, a mutational signature that reflects a defect in DNA double strand break (DSB) repair (Supplemental Figure 1B-C)(19). In conjunction with previously-published work (9, 12, 13, 20) these findings motivated us to generate a mouse model of *Atrx* deleted sarcoma and examine the response to DNA DSB-inducing therapeutics.

**Figure 1:**
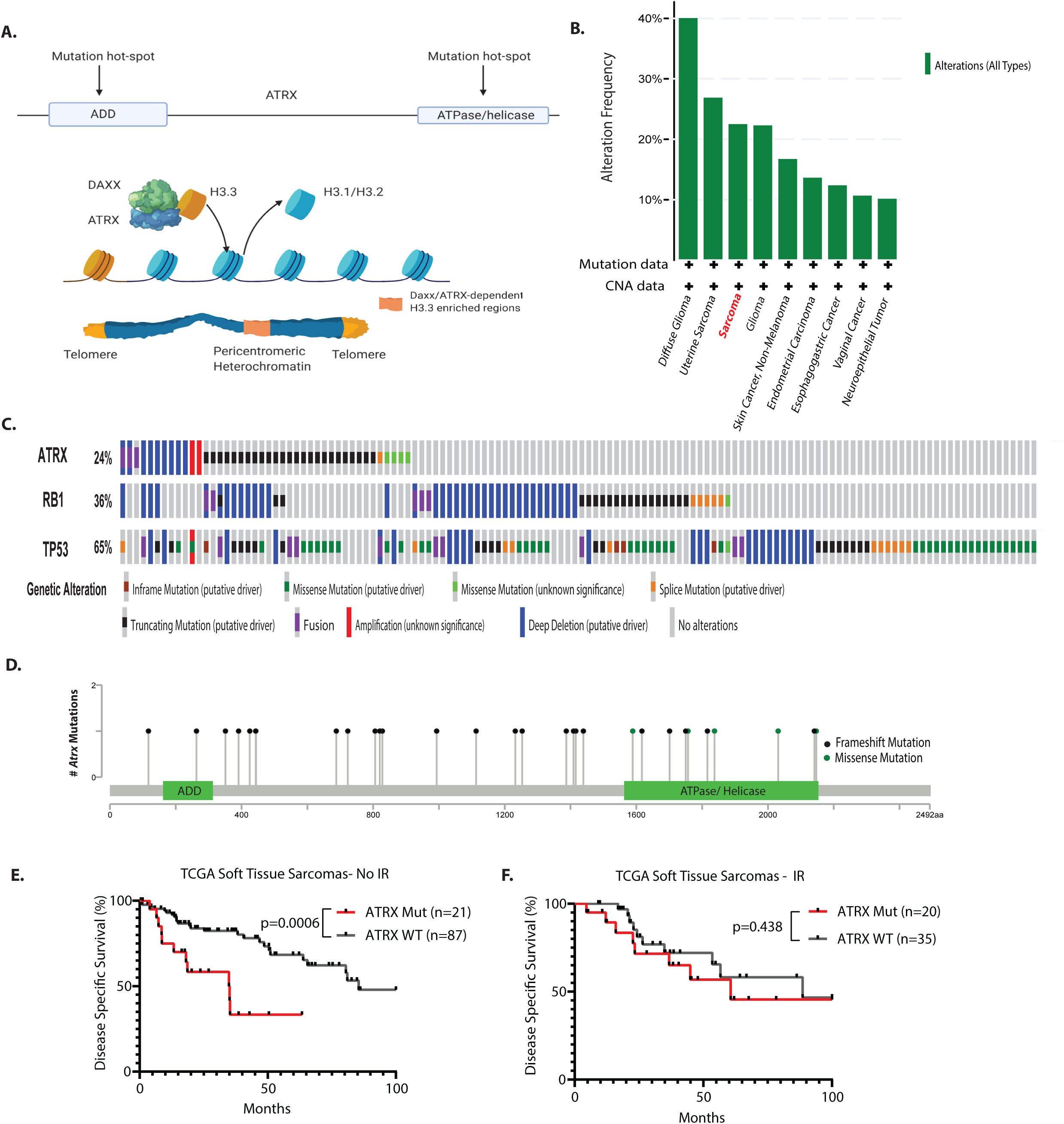
Genomic characteristics of ATRX in human cancer. A) Diagram with two major functional domains of ATRX (ADD and ATPase/helicase). ATRX complexes with DAXX to promote deposition of histone 3.3 throughout the genome, with especially high concentrations at constitutive heterochromatin. B) Alterations in *ATRX* in human cancers from the TCGA sequencing database. C) Schematic of the most frequently mutated genesin a human soft tissue sarcoma cohort comprised of leiomyosarcoma, myxofibrosarcoma, and undifferentiated pleomorphic sarcoma (STS cohort), in the TCGA sequencing database with each vertical row representing a single tumor sample. D) Positional distribution of mutations in the *ATRX* gene from the STS cohort from the TCGA. Black lollipop markers represent insertion or deletion mutations, while green lollipop markers represent missense mutations. E) Disease specific survival for an unirradiated STS cohort, comparing tumors with ATRX genomic alterations (n=21) and tumors with wild-type ATRX (n=87). Statistical comparison was performed using a Log-rank (Mantel-Cox) test. F) Disease specific survival for a STS cohort in which tumors received radiation therapy, comparing tumors with ATRX genomic alterations (n=20) and tumors with wild-type ATRX (n=35). Statistical comparison was performed using a log-rank (Mantel-Cox) test.

### Generation and characterization of a primary mouse model of soft tissue sarcoma with *Atrx* deletion

To study the effect of *Atrx* in soft tissue sarcoma, we adapted a spatially and temporally restricted primary mouse model of soft tissue sarcoma (P7 KPA model) (21). This model utilizes an estrogen receptor-regulated Cre-recombinase that is expressed from the muscle satellite cell-specific *Pax7* promoter (Pax7-CreER^T2^). The model also contains a conditional oncogenic *Kras^G12D^* allele that is preceded by a stop cassette flanked by loxp sites (floxed) at the endogenous *Kras* promoter (*LSL-Kras^G12D^*). Finally the model has two copies of *Trp53* with floxed exons 2 through 10 (*p53^FL^*), and *Atrx* allele(s) that with a floxed exon 18, which is required for SWI/SNF protein function (*Atrx^FL^*). Because *Atrx* is X-linked, two *Atrx^FL^* alleles are required in female mice and one *Atrx^FL^* allele is required in male mice to generate tumors lacking *Atrx* expression. To initiate a sarcoma in the P7 KPA model (Figure 2A), we injected mice in the gastrocnemius muscle with 4-hydroxytamoxifen (4-OHT) to activate Cre recombinase in muscle satellite cells. Once activated, Cre drives excision of the stop cassette preceding oncogenic Kras*^G12D^* and leads to deletion of *Trp53* and *Atrx* floxed alleles (Figure 2A). For all experiments, we used littermate control mice that retained at least one wild-type *Atrx* allele (P7 KP model, Figure 2A). Time to tumor detection following intramuscular 4-OHT injection was delayed in P7 KPA mice compared to P7 KP mice that retained *Atrx* expression: P7 KPA median 55 days (range 27-77) vs. P7 KP median 35 days (range 27-61). The histologic appearance of P7 KPA sarcomas stained with hematoxylin and eosin (H&E) or myogenic markers was similar to P7 KP sarcomas, with most of the tumors mimicking human UPS (Figure 2B, Supplementary Figure 2) as previously described (21). A minority of the sarcomas in the mouse P7 KPA model were classified as rhabdomyosarcoma due to the presence of rhabdomyoblasts and myogenic markers (Figure 2C). There was a similar distribution of sarcoma sub-types identified within the P7 KP control cohort (Figure 2D). To assess *Atrx* recombination in the P7K KPA model, we generated cell lines from primary tumors. After culturing the cells *in vitro* for at least eight passages to eliminate stromal cell contamination, we isolated genomic DNA and performed PCR genotyping for *Atrx*. As expected, the loxP flanked exon 18 of Atrx was efficiently recombined in the P7 KPA model (Figure 2E).

**Figure 2.**
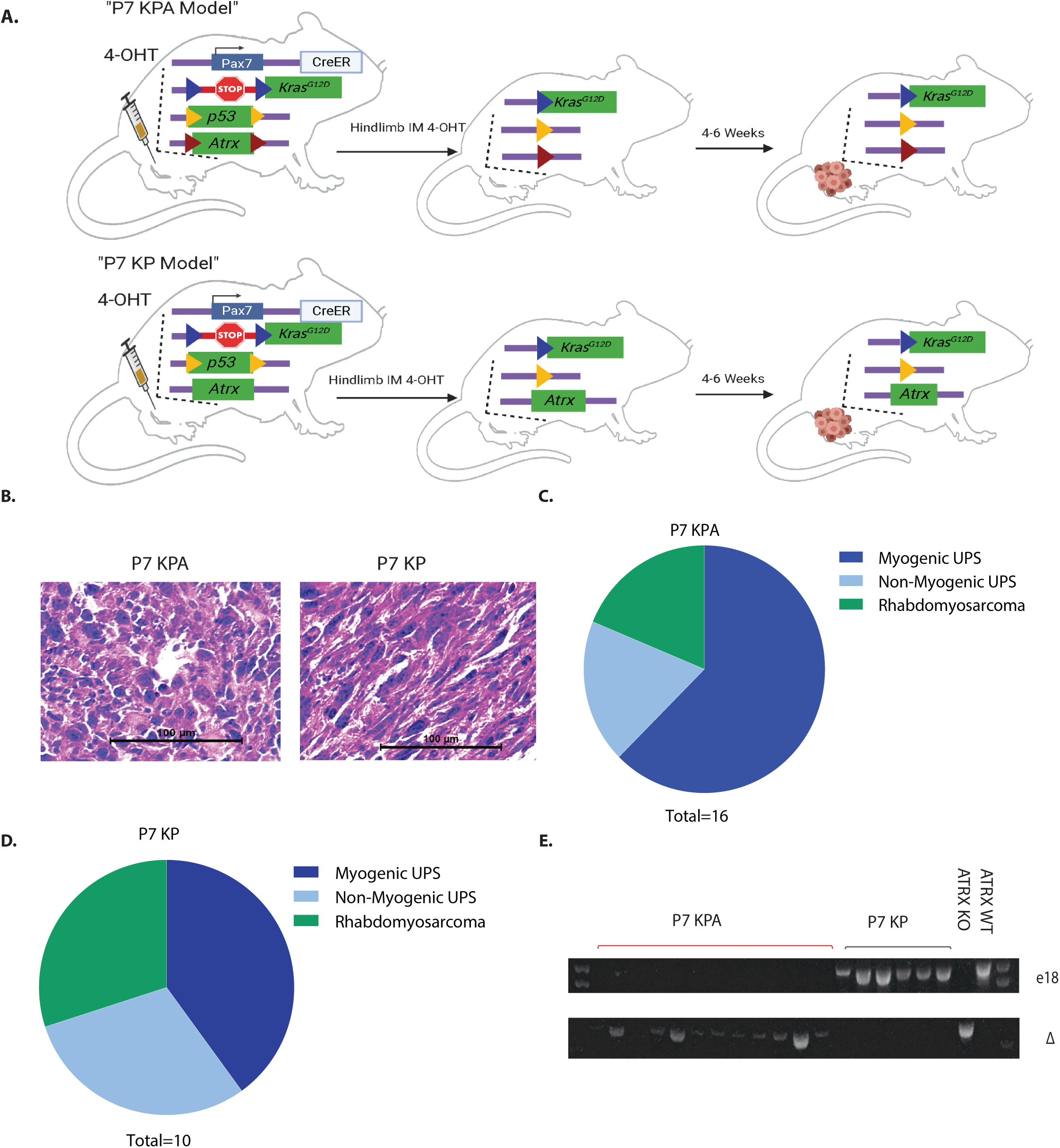
Primary mouse model of *Atrx* deleted soft tissue sarcoma. **A)** Schematic showing the spatially and temporally restricted primary mouse model of *Atrx* deleted soft tissue sarcoma. 4-hydroxytamoxifen (4-OHT) is injected into the gastrocnemius muscle, which leads to activation of the CreER expressed from the endogenous Pax7 promoter in muscle satellite cells to activate expression of Kras^G12D^ and delete floxed p53 and *Atrx* alleles. *Atrx* is deleted in P7 KPA mice (top) and a wild type *Atrx* is retained in control P7 KP mice (bottom). Loxp sites are designated by colored triangles in the diagram. **B)** Hematoxylin and eosin (H&E) staining of sarcomas from P7 KPA and P7 KP mice. These H&E images are also shown in Figure 1A (left) accompanied by staining for myogenic markers. **C-D)** Classification of tumor type, as determined by IHC for 9 myogenic and other markers. Myogenic undifferentiated pleomorphic sarcoma (UPS) was defined as when cells had pleomorphic nuclei characteristic of UPS but stained positive for at least 2 of the four tested myogenic markers (MyoD1, Myogenin, Desmin and SMA). **E)** Genotyping of sarcoma cell lines from P7 KP and P7KPA mouse tumors for the presence of loxp flanked *Atrx* exon 18 (top) and recombined *Atrx* with deletion of exon 18 (bottom).

### *Atrx* deletion increases sensitivity to DNA double strand break-inducing therapies *in vitro*

To test how *Atrx* deletion impacted the chemotherapy response *in vitro*, we first generated murine UPS cell lines with activated oncogenic *Kras^G12D^* and *p53* deletion from KP mouse model sarcomas (22). We then used CRISPR-Cas9 gene editing to delete *Atrx* and used a vector only control in this same cell line to generate an isogenic *Atrx* wild type control. We confirmed successful knockout of *Atrx* in the ATRX KO isogenic lines at the protein level (Figure 3A-B). Next, we tested whether *Atrx* deletion increased sensitivity to the DNA strand break inducing chemotherapy doxorubicin (Figure 3C). Since we observed an increased sensitivity to double strand break-inducing therapy, we reasoned that deletion of *Atrx* may also increase sensitivity to ionizing radiation in *vitro*. We performed clonogenic assays using single radiation doses (2 Gy to 8 Gy) and observed that *Atrx* deletion decreased sarcoma cell survival following irradiation in three different isogenic primary cell line pairs (Figure 3D-F).

**Figure 3:**
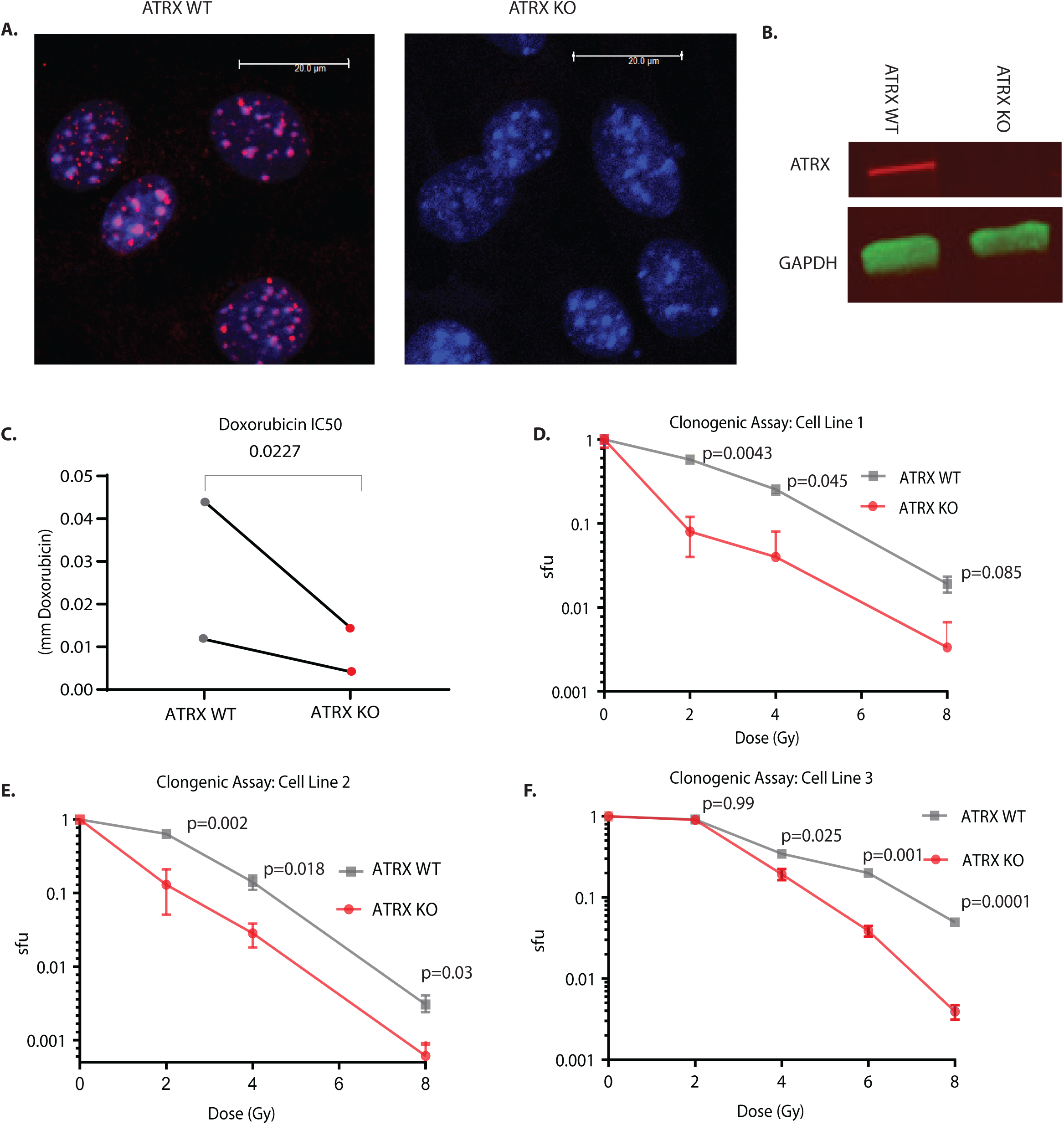
*Atrx* deletion increases sensitivity to DNA double strand break-inducing therapeutics. A) Immunofluorescence staining for ATRX (red) in isogenic cell lines derived from single cells after transfection with Cas9 and either a vector only control(left, “ATRX WT”), or a sgRNA to ATRX (right, “ATRX KO”) or. B) Western blot of ATRX and GAPDH in ATRX WT and ATRX KO cell lines. C) IC50 assays in which cell lines were treated with the DNA double strand break inducing therapeutic doxorubicin. Statistical analysis was performed using a ratio paired t-test. Each data point represents a biological replicate. D-F) Clonogenic assay of isogenic *Atrx* deleted and intact cell line pairs after the indicated doses of ionizing radiation. Surviving fraction (sfu) is shown in log scale on the y-axis. Statistical analysis was performed using multiple Welch’s t-tests corrected for multiple comparisons using the Holm-Šídák method. Each graph represents a separate biological replicate isogenic cell line pair.

### *Atrx* deletion results in persistent DNA damage, telomere dysfunction-induced foci, and severe mitotic defects after irradiation

Next, we investigated the mechanism by which loss of *Atrx* increased radiosensitivity in our soft tissue sarcoma cell lines. Because of the DNA damage associated signature we detected in *ATRX* mutant human UPS (Supplementary Figure 1B-C) and because of prior reports describing a role for ATRX in DNA damage and telomere protection (13, 20), we hypothesized that deletion of *Atrx* would reduce DNA damage repair efficiency at telomeres, causing an increase in persistent DNA double strand breaks in these regions after irradiation. Persistent telomere breaks can lead to genomic instability, mitotic dysfunction, and cell death (23, 24). To test this hypothesis, we performed immunofluorescence staining for DNA double strand break protein 53BP1 in conjunction with telomere fluorescence *in situ* hybridization (immunoFISH) in paired *Atrx* WT and *Atrx* KO isogenic cell lines three days after treatment with 4 Gy (Figure 4A-F, Supplementary Figure 3). This immunoFISH experiment demonstrated that *Atrx* deletion significantly increased 53BP1 foci 3 days after irradiation, suggesting a global increase in persistent DNA damage (Figure 4B). Additional analysis revealed that *Atrx* deletion increased the number of 53BP1 foci that co-localized with telomeres, also known as telomere dysfunction induced foci (TIF), suggesting that functional ATRX protects telomeres in sarcoma cells from radiation-induced DNA damage and TIFs (Figure 5A, C).

**Figure 4:**
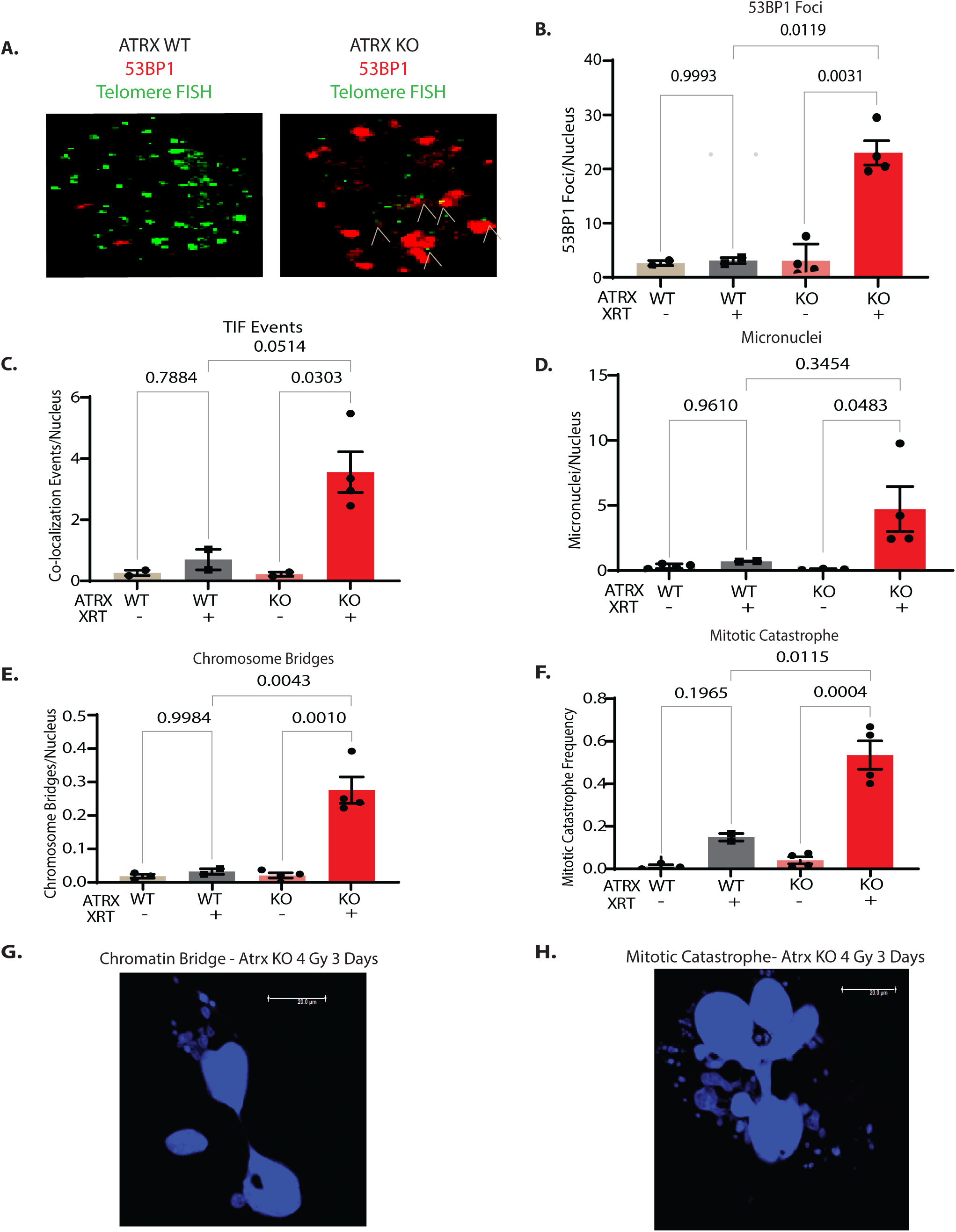
*Atrx* deletion leads to persistent DNA double stranded breaks and telomere dysfunction after irradiation. A) Representative images showing 53BP1 Foci (red) and telomere foci (green) in ATRX WT (left) and ATRX KO (right) cell lines three days after 4 Gy. White markers show colocalization of telomere fish foci and 53 BP1 foci, which are telomere dysfunction induced foci (TIF). B-F) Quantification of an *Atrx* isogenic cell line pair assayed three days after 4 Gy. Each dot represents an experimental repeat immunoFISH experiment of a single isogenic cell line pair with at least 7 images quantified for each experiment. Replicates are: ATRX KO + IR (n=4), ATRX KO untreated (n=4), ATRX WT + IR (n=2), ATRX WT untreated (n=3). Data are shown for 53BP1 foci (B), 53BP1 and telomere FISH co-localization marking TIFs (C), micronuclei (D), persistent chromosomal bridges between cells (E), and mitotic catastrophe (F). Statistical analysis was performed using a two-way ANOVA with Tukey’s multiple comparisons test (B-F). G) DAPI staining showing a chromatin bridge in an *Atrx* KO cell line treated with 4 Gy and assayed three days later. H) DAPI staining showing a cell undergoing mitotic catastrophe in an *Atrx* KO cell line treated with 4 Gy and assayed three days later.

**Figure 5.**
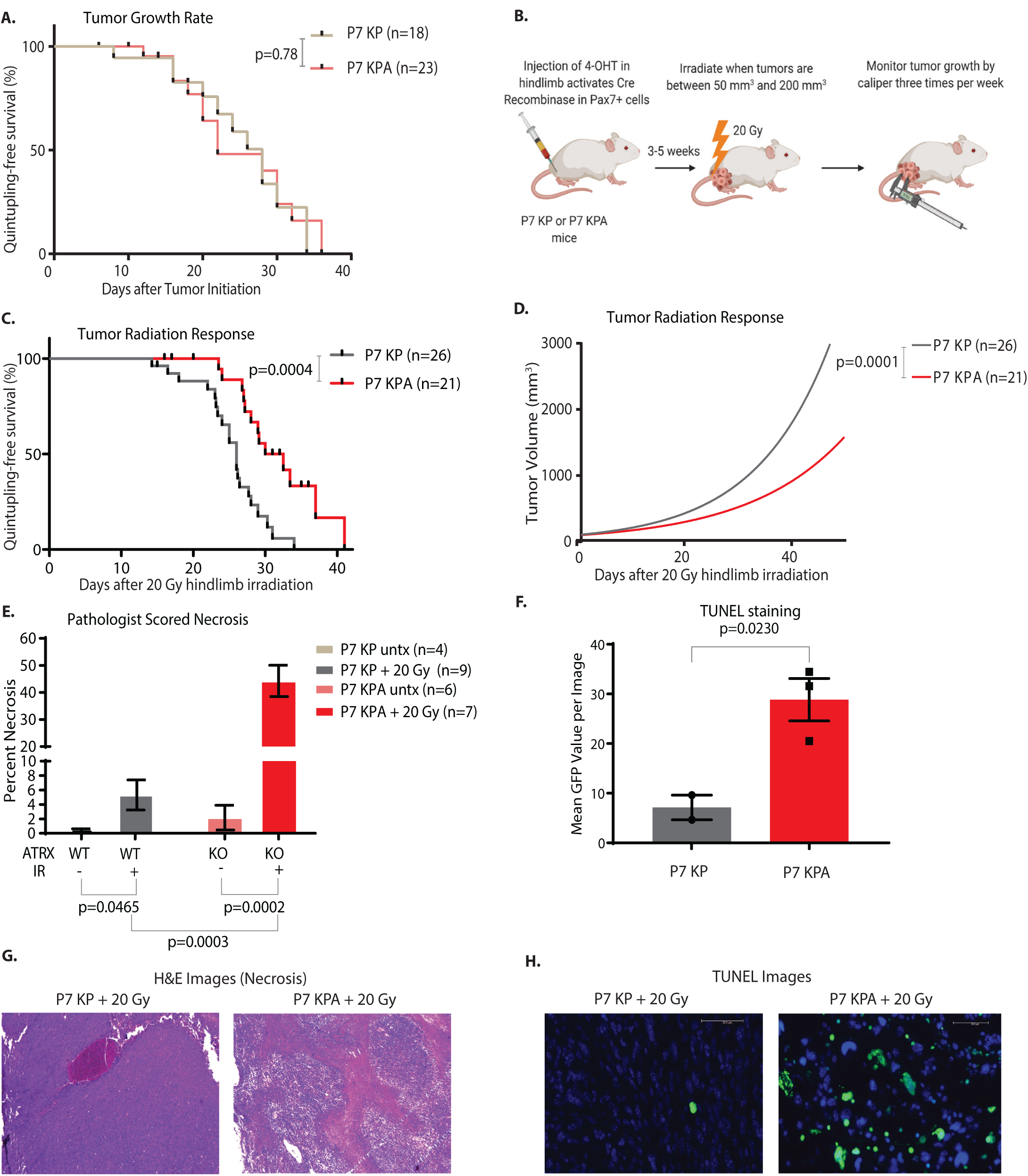
*Atrx* deletion increases radiosensitivity *in vivo*. A) Tumor growth rates of sarcomas that retained *Atrx* in P7 KP mice and sarcomas that deleted *Atrx* in P7 KPA mice as measured by time for tumor to quintuple in size relative to size at initial measurement. Comparison of survival curves was performed using log-rank (Mantel Cox) test. B) Diagram summarizing experimental procedure for mouse sarcoma irradiation and monitoring. C) Measurement of tumor growth rates after 20 Gy hindlimb irradiation, as measured by time for tumor to quintuple in size relative to size at treatment. Comparison of survival curves was performed using log-rank (Mantel Cox) test. D) Non-linear fit modeling of tumor growth curves after 20 Gy hindlimb irradiation. E) Quantification of pathologist scored percent area necrosis of samples that were either untreated (lighter colors) or treated with 20 Gy (darker colors) and harvested at 6 days. Each data point represents a biological replicate. F) Quantification of TUNEL positive fluorescence in frozen sections of tumor samples harvested 3 days after 20 Gy of hindlimb radiation. Each data point represents a biological replicate. G) Representative H&E staining from sarcomas in P7 KP (left) and P7 KPA (right) mice harvested 6 days after 20 Gy. Images were scored for necrosis by a sarcoma pathologist. H) Representative TUNEL staining from sarcomas from P7 KP (left) and P7 KPA (right) mice harvested 3 days after 20 Gy. GFP positive cells are TUNEL positive.

As persistent DNA double strand breaks can lead to chromosome mis-segregation and mitotic error, we next evaluated whether irradiation of sarcoma cells with *Atrx* deletion caused increased mitotic dysfunction, including micronucleus formation, chromosome bridges, and mitotic catastrophe events. Quantification showed that the *Atrx* KO cell line had a significant increase in micronuclei after radiation, but we did not detect enhanced micronucleus formation in the *Atrx* WT cell line (Figure 4D). Irradiation of the *Atrx* deleted sarcoma cell line also significantly increased chromosome bridge events and the number of cells undergoing mitotic catastrophe relative to its irradiated *Atrx* WT counterpart (Figure 4E-H). Consistent with the enhanced mitotic dysfunction, immunofluorescence staining of the nuclear envelope protein Lamin B1 with DAPI counterstain showed that *Atrx* deletion increased micronuclei after treatment with ionizing radiation across multiple isogenic cell line pairs (Supplementary Figure 5D-E, Figure 7A).

### *Atrx* deletion radiosensitizes and increases cell death in sarcomas in autochthonous primary mouse models

Next, we set out to evaluate whether *Atrx* deletion increases tumor response to radiation therapy in the primary P7 KP mouse model. Loss of *Atrx* had no discernable effect on sarcoma growth rates in the absence of treatment, but following a single dose of 20 Gy focal radiation therapy tumors in P7 KPA mice demonstrated a significant growth delay relative to tumors in the P7 KP mice (Figure 5A-D). In a separate cohort of these mice, we harvested sarcomas in P7 KPA and P7 KP mice 6 days after a single fraction of 20 Gy focal irradiation. Examination of hematoxylin and eosin (H&E) stained sections from these tumors revealed that *Atrx* deletion resulted in a significant increase in tumor necrosis 6 days after irradiation (Figure 5E, 5G). In addition, deletion of *Atrx* in sarcomas harvested 3 days post 20 Gy focal irradiation led to a significant increase in cell death detected through terminal deoxynucleotidyl transferase dUTP nick end labeling (TUNEL) staining (Figure 5F, 5H). P7 KPA tumors also had a significantly lower fraction of Ki67 positive cells by immunohistochemistry (IHC) 6 days after radiation therapy (Supplementary Figure 4A-B). To evaluate whether this phenotype was limited to the *Kras^G12D^* expressing P7 KP model, we then repeated this experiment in a second primary mouse model of soft tissue sarcoma. The P7 P + MCA model does not utilize a genetically engineered conditional oncogenic *Kras^G12D^* allele, but instead is initiated by intramuscular injection of 4-OHT in *Pax7-CreER^T2^*; *p53^FL/FL^* mice to delete *p53*, followed by injection of 3-methylcholanthrene (MCA), a potent carcinogen that drives base substitutions at the site of injection (Supplementary Figure 5A) (25). Similar to the P7KPA model, *Atrx* deletion in the P7 P + MCA model significantly delayed tumor growth after radiation as measured by time to tumor quintupling (Supplementary Figure 4C). Therefore, deletion of *Atrx* causes radiosensitivity in two independent primary mouse models of soft tissue sarcoma.

### ATRX deleted sarcomas have reduced adaptive and innate immune signaling after ionizing radiation

We then evaluated whether the observed increase in radiosensitivity *in vivo* after *Atrx* deletion was mediated by the adaptive immune system. To test this, we first performed RNA-sequencing of cohorts of P7 KPA and P7 KP sarcomas that were not irradiated. Both gene ontology (GO) and reactome pathway enrichment analysis of these samples revealed a marked downregulation of adaptive immune signaling in the P7 KPA cohort compared to the P7 KP control (Figure 6A-B). We then performed RNA sequencing of P7 KPA and P7 KP tumors harvested six days after 20 Gy of radiation therapy. GO and reactome pathway enrichment analysis of these tumors also demonstrated a marked downregulation of adaptive immune signaling. Both “interferon gamma response” (GO) and “immunoregulatory interactions between a lymphoid and non-lymphoid cell” (reactome) were the second most highly enriched pathways in our analysis of downregulated gene sets in irradiated P7 KPA tumors compared to the irradiated P7 KP controls (Figure 6C-D). We then estimated the level of T-cell and macrophage/monocyte infiltration in the irradiated P7 KPA and P7 KP tumors using mMCP counter analysis of the RNA-sequencing data (26). While these comparisons failed to demonstrate a significant difference, there was a trend towards lower levels of both T-cell and macrophage infiltrates in the irradiated P7 KPA tumors when compared to the irradiated P7 KP counterparts. Taken together, these data indicate that *Atrx* deletion decreases adaptive immune signaling after radiation therapy in soft tissue sarcomas.

**Figure 6.**
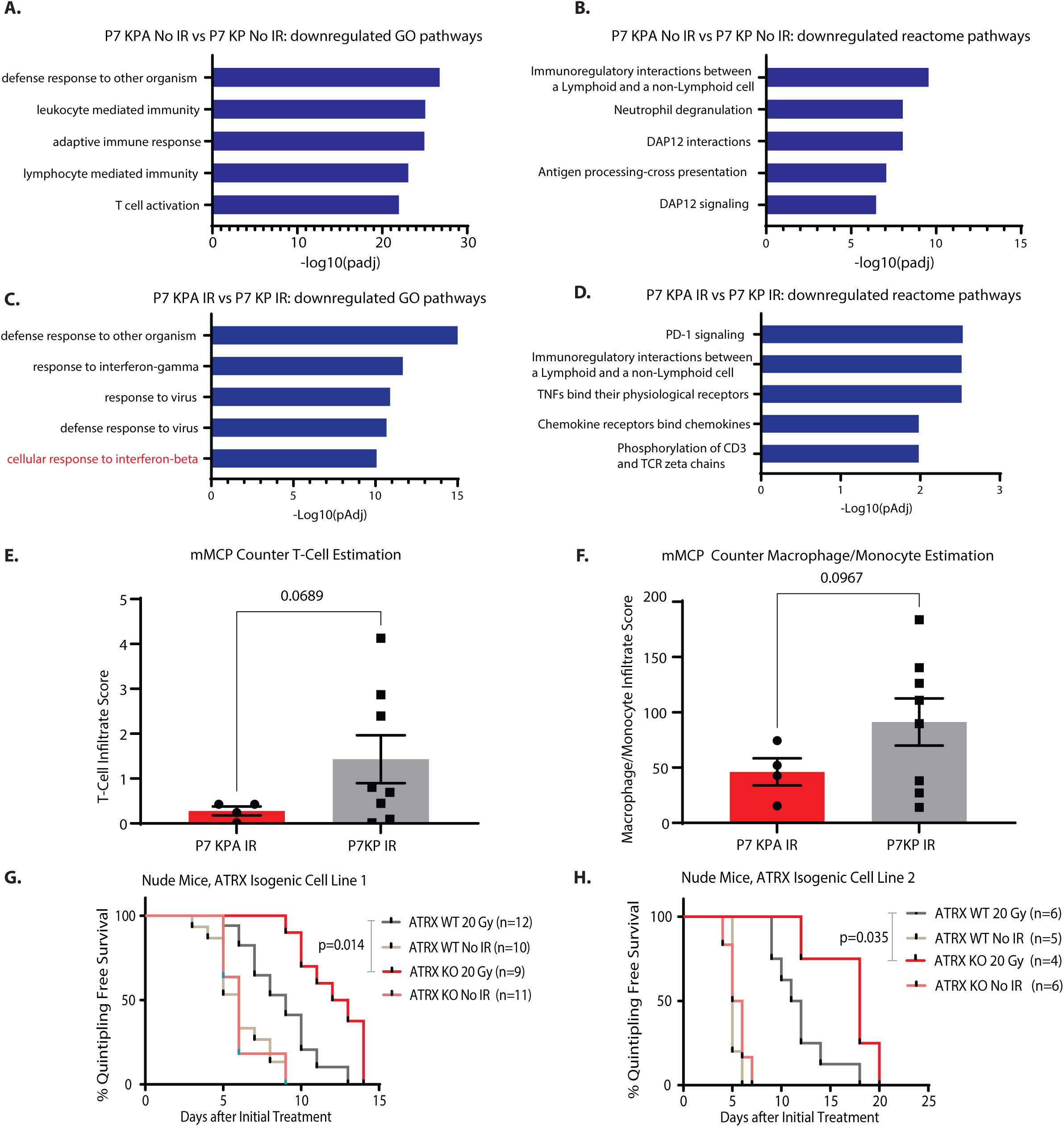
ATRX deletion radiosensitizes STS in a T-cell independent manner. A) Top five results for RNA sequencing GO pathway enrichment analysis comparing unirradiated P7 KPA sarcomas (n=4) vs unirradiated P7 KP sarcomas (n=4). B) Top five results for RNA sequencing reactome pathway enrichment analysis comparing unirradiated P7 KPA sarcomas (n=4) vs unirradiated P7 KP sarcomas (n=4). C) Top five results for RNA sequencing GO pathway enrichment analysis comparing P7 KPA sarcomas harvested six days after treatment with 20 Gy ionizing radiation (n=4) vs P7 KP sarcomas harvested six days after treatment with 20 Gy ionizing radiation (n=8). D) Top five results for RNA sequencing reactome pathway enrichment analysis comparing P7 KPA sarcomas harvested six days after treatment with 20 Gy ionizing raidation (n=4) vs P7 KP sarcomas harvested six days after treatment with 20 Gy ionizing radiation (n=8). E) Estimation of T-Cell infiltrate via mMCP counter analysis of RNA sequencing data, comparing P7 KPA sarcomas harvested six days after treatment with 20 Gy ionizing radiation (n=4) to P7 KP sarcomas harvested six days after treatment with 20 Gy ionizing radiation (n=8). Each data point represents a biological replicate. F) Estimation of Macrophage/Monocyte infiltrate via mMCP counter analysis of RNA sequencing data, comparing P7 KPA sarcomas harvested six days after treatment with 20 Gy ionizing radiation (n=4) to P7 KP sarcomas harvested six days after treatment with 20 Gy ionizing radiation (n=8). Each data point represents a biological replicate. G-H) A single cell line from one of two ATRX isogenic cell line pairs was implanted into the hindlimb muscle of nude mice. Once tumors formed, mice were randomized to receive radiation therapy or no treatment. Survival curves were estimated for each group, considered separately, using the Kaplan-Meier method and compared statistically using the log-rank (Mantel Cox) test.

### *Atrx* deletion sensitizes sarcomas to radiation independently of T-cell mediated immunity

Based on these findings, we then investigated whether the deletion of ATRX could radiosensitize soft tissue sarcoma in a T-cell independent manner. We transplanted isogenic KP sarcoma cell lines with and without *Atrx* deletion into the hindlimb muscles of athymic nude mice and measured the rate of tumor (Supplementary Figure 4B). We found that *Atrx* deletion radiosensitized the transplanted tumors even in the absence of an intact immune system (Figure 6G-H). Together, these results showed that *Atrx* deletion increases radiosensitivity in both T-cell-deficient and immunocompetent models *in vivo*.

### *Atrx* deletion impairs cGAS-STING mediated type-I interferon response after radiation

When comparing irradiated P7 KPA and P7KP tumors, we noted that in addition to downregulation of adaptive immune signaling there was also a significant downregulation of antiviral defense and type-I interferon signaling related pathways (Figure 6C). In the context of radiation therapy, one prominent mechanism for activation of type-I interferon signaling is the cGAS-STING pathway. Micronuclei can rupture, releasing double strand DNA into the cytoplasm that activates the cytoplasmic double strand DNA sensing system cGAS-STING to drive expression of type-I interferon (17). Because *Atrx*-deleted sarcoma cells had increased micronuclei after irradiation (Figure 7A), we next tested whether *Atrx* deletion increased interferon beta (*Ifnb1*) expression after ionizing radiation *in vitro*. Interestingly, we found that *Atrx* deletion reduced *Ifnb1* expression following 4 Gy relative to the matched isogenic *Atrx* WT control (Figure 7B). To determine whether this finding was the result of an impairment in the cGAS-STING pathway, we first assessed the proficiency of the pathway in mouse sarcoma cell lines via cGAS-STING induction using interferon stimulatory DNA (ISD). ISD is 45 base pair non-CpG oligomer derived from the *Listeria monocytogenes* genome that is known to induce cGAS-STING activation (27). We found that after transfections with ISD, cells with *Atrx* deletion had markedly decreased cGAS-STING mediated *Ifnb1* expression (Figure 7C). To test whether impairment of cGAS-STING would reduce radiation-induced type I interferon signaling in *Atrx* deleted sarcomas *in vivo*, we collected tumors six days after treatment with 20 Gy from P7 KP or P7 KPA mice. After irradiation *in vivo*, we observed that *Atrx* deletion also impaired type-I interferon signaling (Figure 7D). Therefore, we performed RNA sequencing on three ATRX intact and deficient isogenic sarcoma cell line pairs after treatment with either ISD, 4 Gy, or untreated controls. Pathway analysis revealed a significant downregulation of type-I interferon signaling In *Atrx* deficient cell lines relative to their ATRX intact controls after treatment with ISD (Figure 7E) or ionizing radiation (Figure 7F). To determine what part of the cGAS-STING signaling pathway was impaired following deletion of *Atrx*, we next treated the isogenic cell lines with DMXAA, a STING agonist. DMXAA treatment induced significantly less *Ifnb1* expression in *Atrx* deleted cell lines (Figure 7G), suggesting that the impairment in cGAS-STING signaling was occurring downstream of cGAS or cGAMP. An alternative explanation for these results is that loss of ATRX function disrupts *Ifnb1* gene transcription independent of cGAS-STING signaling. Therefore, we treated cell lines with and without *Atrx* deletion with Poly(I:C), an RNA-based compound known to stimulate *Ifnb1* via a cGAS-STING independent pathway. Stimulation with Poly(I:C) induced greater than 25-fold increase in *Infb1* expression relative to untreated controls in every *Atrx* intact and deleted cell line tested (Figure 7H). Together these data demonstrate that *Atrx* deletion impairs response to the cGAS-STING pathway in soft tissue sarcoma downstream of cGAS signaling but upstream of *Infb1*.

**Figure 7:**
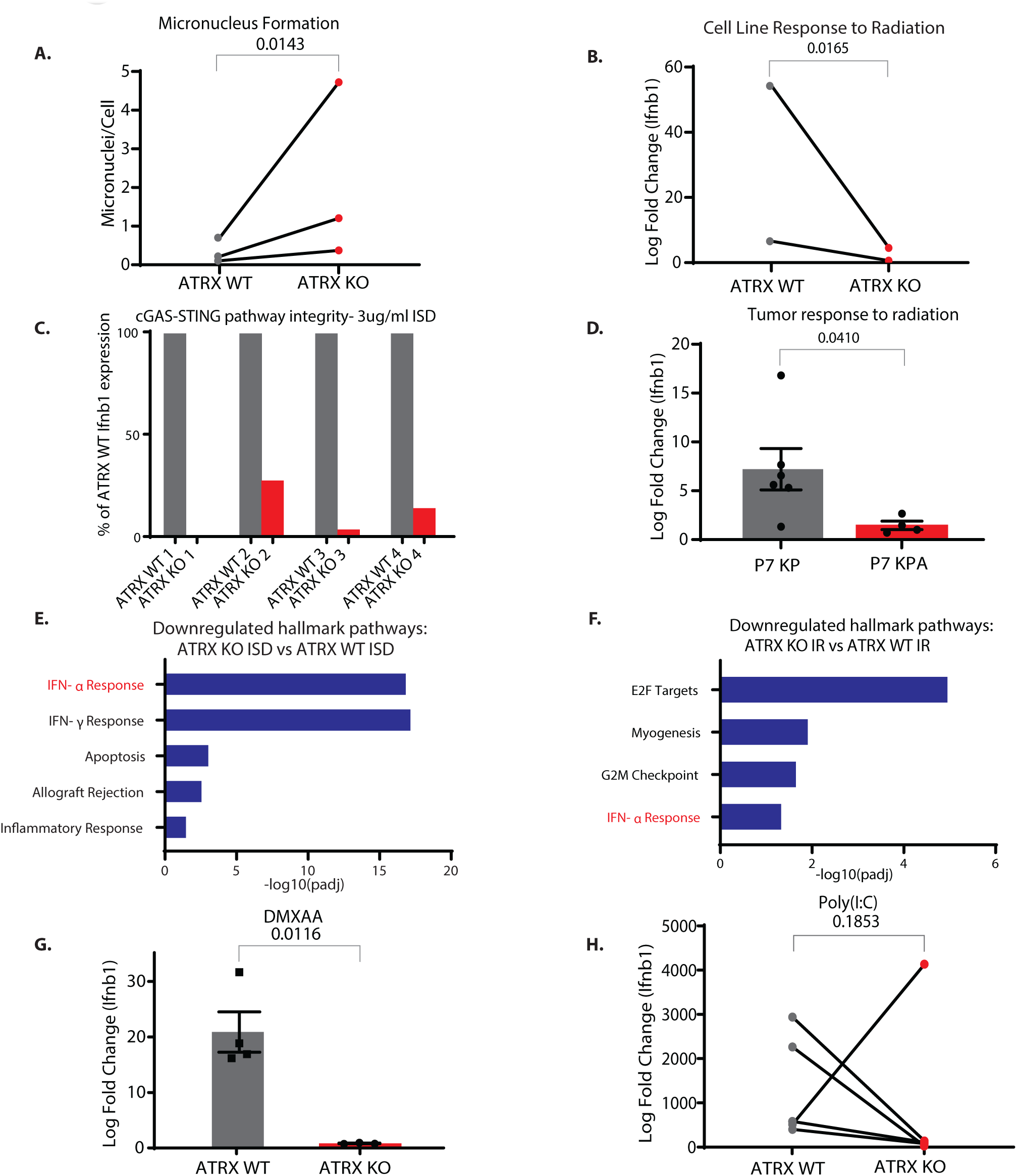
*Atrx* deletion suppresses type-I interferon signaling in a cGAS-STING-dependent manner. A) Quantification of micronuclei for cell lines analyzed three days after treatment with 4 Gy. Each dot represents a biological replicate, and each biological replicate had at least five separate fields scored. For statistical analysis, a ratio paired T-test was performed, pairing each *Atrx* KO cell line to its *Atrx* WT isogenic cell line counterpart. B) RT-PCR quantification of *Ifnb1* in the indicated isogenic sarcoma cell line harvested 24 hours after transfection with interferon stimulatory DNA (ISD). Each *Atrx* KO isogenic cell line has been normalized to its *Atrx* WT counterpart to enable visualization and comparison between biological replicates. C) RT-PCR quantification of log fold change expression of *Ifnb1* of isogenic cell lines assayed three days after 4 Gy of ionizing radiation. For statistical analysis, a ratio paired T-test was performed, pairing each *Atrx* KO cell line to its *Atrx* WT isogenic cell line counterpart. D) RT-PCR quantification of log fold change expression of *Ifnb1* of sarcomas from P7 KP and P7 KPA mice harvested 6 days after treatment with 20 Gy of ionizing radiation relative to its untreated control. Statistical comparison utilized an unpaired t-test with Welch’s correction. E) Hallmark pathway gene set enrichment analysis of RNA sequencing data comparing 3 ATRX KO isogenic cell lines treated with ISD to 3 ATRX WT isogenic cell lines treated with ISD. F) Hallmark pathway gene set enrichment analysis of RNA sequencing data comparing 3 ATRX KO isogenic cell lines treated with 4 Gy and harvested 36 hours after treatment to 3 ATRX WT isogenic cell lines treated with 4 Gy and harvested 36 hours after treatment. G) RT-PCR of *Ifnb1* for isogenic cell line pair 1 treated for 24 hours with 100 ug/ml DMXAA, a potent STING agonist. Each dot represents a separate experimental repeat of the RT-PCR assay. Statistical analysis was performed using an unpaired t-test with Welch’s correction H) RT-PCR of *Ifnb1* for ATRX isogenic cell line pairs treated with Poly(I:C) at a concentration of 1 ug/ml and harvested at 24 hours. Statistical analysis was performed using a ratio paired t-test. All data points represent biological replicates.

### Deletion of *Atrx* increases sensitivity to oncolytic herpesvirus therapy

Activation of cGAS-STING in response to viral dsDNA leads to expression of type I interferon inflammatory cytokines, which limits viral replication and spread of herpes simplex virus type 1 (HSV-1) and herpesvirus based oncolytic viral cancer therapy (Figure 8A) (27). KEGG pathway enrichment analysis of RNA-sequencing data revealed a marked downregulation of the HSV-1 response pathway in P7 KPA tumors relative to P7 KP controls (Supplemental Figure 6A-B). Based on this downregulation and the finding that *Atrx* deletion reduced activation of the cGAS-STING response to microbial-derived dsDNA, we investigated whether *Atrx* deletion increased sensitivity to an oncolytic HSV-1 designed to specifically target and kill tumor cells. We first tested the effect of transfection of dsDNA from HSV type I (HSV-60), a well described stimulant of cGAS-STING, in three isogenic sarcoma cell line pairs. There was significantly less induction of type-I interferon expression in the ATRX KO cell lines after HSV-60 stimulation when compared to the ATRX WT cell lines (Figure 8B). We therefore tested the impact of *Atrx* deletion on sarcoma cell infection with the oncolytic herpesvirus oHSV, which is a variant of HSV-1 that has been modified to specifically target and destroy tumor cells (28). In 2015, the FDA approved a related oncolytic herpesvirus for the treatment of advanced melanoma (29). Other variants, including the G207 oHSV which is closely related to the virus used for this study, are currently in phase 2 clinical trials for the treatment of recurrent supratentorial brain tumors (NCT03911388, NCT02457845). To assess whether deletion of *Atrx* sensitized sarcoma cells to oHSV, we performed an IC50 assay using four different isogenic cell line pairs. We found that *Atrx* deletion significantly increased sensitivity to oHSV in *Atrx* deficient sarcoma cell lines for every isogenic pair tested (Figure 8C). To extend these data, we performed clonogenic assays, in which we treated both the ATRX WT and ATRX KO cell lines with an identical concentration of oHSV. We observed a significant reduction in the number of colonies formed in the ATRX KO cell lines (Figure 8D-E). Together, these findings demonstrate that ATRX deletion impairs cGAS-STING response to HSV-1 dsDNA and increases susceptibility to oncolytic herpesvirus therapy in soft tissue sarcoma.

**Figure 8:**
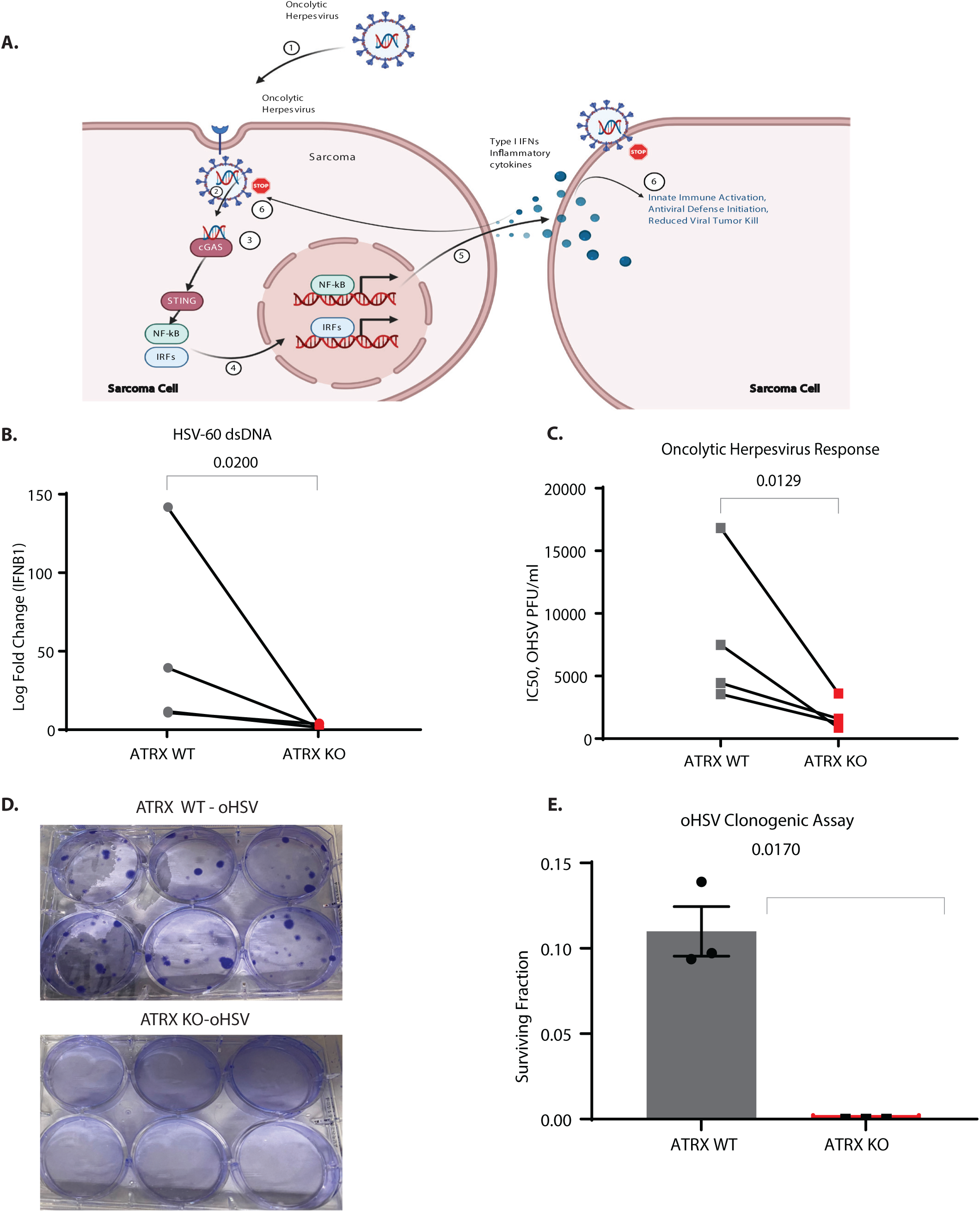
*Atrx* deletion increases sarcoma sensitivity to oncolytic herpesvirus therapy. A) Diagram showing the interaction between oncolytic herpesvirus and cGAS-STING pathway in ATRX WT sarcoma cells. Oncolytic herpesvirus dsDNA is detected and cGAS-STING signaling activates the innate immune response, which inhibits viral spread and oncolytic herpesvirus efficacy. B) RT-PCR of *Ifnb1* for ATRX WT and ATRX KO isogenic sarcoma cell lines assayed 24 hours after treatment with HSV-60, a 60 bp dsDNA oligonucleotide derived from HSV-1 that is a known inducer of cGAS-STING signaling. For statistical analysis, a ratio paired t-test was performed, pairing each ATRX KO cell line to its ATRX WT isogenic cell line counterpart. C) IC50 assays for isogenic sarcoma cell lines treated with oncolytic herpesvirus (oHSV). Differences between all paired cell lines are statistically significant (p<.05) as analyzed by a ratio paired t-test. Each data point represents a biological replicate. D) ATRX WT and ATRX KO paired isogenic cell lines were plated in triplicate at identical density and treated with oHSV at a concentration corresponding to the IC90 of the ATRX WT cell line. Colonies were stained with crystal violet. Each data point represents a biological replicate. E) Quantification of colonies in *Atrx* retained and *Atrx* deleted isogenic cell line pair after oHSV treatment. Each dot represents an experimental repeat using the same cell line pair. Statistical comparison was performed using an unpaired t-test with Welch’s correction.

## Discussion

The contribution of ATRX to tumor development and treatment response is not well defined, despite mutations in *ATRX* occurring in approximately 6% of all human cancers (4). Using a mouse model of soft tissue sarcoma, we identified two categories of therapeutic vulnerability in tumors lacking *Atrx* function. Specifically, we found that *Atrx* loss of function mutations in soft tissue sarcoma confer sensitivity to established therapies, such as radiation therapy, as well as emerging therapies, such as oncolytic herpesvirus.

Here, we developed a primary mouse model that recapitulates some of the most frequently occurring hallmarks of human undifferentiated pleomorphic soft tissue sarcoma: loss of p53 function, upregulated RAS pathway signaling, and loss of *Atr*x (4, 30, 31). This sarcoma model has the advantages that tumors arise in a spatially- and temporally-restricted manner and in a native microenvironment with an intact immune system. Our genetic experiments demonstrated that loss of *Atrx* increased radiosensitivity in four model systems: sarcoma cells *in vitro*, sarcoma cells transplanted into immunodeficient mice *in vivo*, primary *Kras^G12D^; p53^-/-^* sarcomas in P7 KP mice, and primary *p53^-/-^* + MCA sarcomas in P7 P mice. Therefore, our findings suggest that *ATRX* mutations may be used to identify a subset of soft tissue sarcomas and potentially other tumors that are more sensitive to radiation therapy in the clinical setting.

Mechanistically, our work indicates that loss of function mutations in *Atrx* drive an increase in persistent DNA damage at telomeres after ionizing radiation therapy. This finding is consistent with previous reports suggesting that ATRX impairs DNA damage repair and increases telomere sister chromatid cohesion defects after induction of telomere double strand breaks (10–13, 32). While this defect was suggested by others to recapitulate key hallmarks of ALT positive cancers, our study also links the increased propensity for TIFs in *Atrx* deleted sarcomas to increased mitotic catastrophe and necrosis after treatment with radiation therapy. Together, these findings define a mechanism for how *Atrx* deletion increases radiosensitivity. We further extended these findings by demonstrating that *Atrx* deletion decreases adaptive immune signaling and interferon signaling after ionizing radiation *in vivo*. We then showed that loss of *Atrx* can radiosensitize soft tissue sarcomas in a T-cell independent manner *in vivo*, as demonstrated by transplant experiments in immunodeficient nude mice.

DNA damage-induced mitotic dysfunction leads to increased micronuclei that frequently rupture, releasing dsDNA into the cytoplasm. This dsDNA can activate cGAS-STING signaling, which thereby increases innate and adaptive immune signaling in tumors (33). The cGAS-STING pathway is emerging as a key regulator of cancer development and therapeutic response with roles in tumor development, metastasis, immunomodulation, senescence, viral defense, and autophagy (34–40). Previous work demonstrated that in a fibroblast cell line knockdown of *Atrx* or its partners *Daxx* and *H3.3* inhibited cGAS-STING signaling in response to high levels of telomeric dsDNA in the cytoplasm of ALT positive cell lines (14). It was unclear, however, whether impaired cGAS-STING signaling would be present in *Atrx*-deleted sarcomas and how radiation would alter this pathway *in vitro* and *in vivo*. Our findings demonstrate that *Atrx* deletion in sarcoma cells impairs cGAS-STING response to both dsDNA transfection *in vitro* and to radiation *in vitro* and *in vivo*. Our results do not exclude the possibility that increased cytoplasmic dsDNA in *Atrx* mutant sarcomas creates a selective pressure for loss of the cGAS-STING signaling pathway. This possibility is less likely, however, because of our experiments with the STING agonist DMXAA and poly (I:C), which revealed a shared dysfunction in cGAS-STING signaling downstream of cGAS, but upstream of *Ifnb1*. Further study is warranted to elucidate the precise manner in which ATRX influences cGAS-STING function. Together these findings provide a mechanistic basis for the effect of ATRX loss of function on type-I interferon signaling. They also provide a rationale for testing treatments in ATRX mutant tumors when the cancer therapy is more effective in the setting of impaired cGAS-STING signaling.

Interestingly, a key function of the cGAS-STING pathway in normal cells is innate immune defense against dsDNA viruses. It has been previously demonstrated that deletion or impairment of the cGAS-STING pathway can increase susceptibility to oncolytic herpesvirus *in vitro* and *in vivo* in both normal cells and cancer (40–45). Our work extends these findings by showing that *Atrx* deletion impairs the cGAS-STING pathway and increases sarcoma susceptibility to oncolytic herpesvirus *in vitro*. ATRX may also contribute to dsDNA antiviral immunity via alterative mechanisms such as maintenance of viral heterochromatin (46). Future studies will be needed to determine whether ATRX-mediated effects on viral heterochromatin also contribute to the sensitivity to oncolytic herpesvirus observed in *Atrx*-deleted sarcoma cells. Regardless of which additional mechanisms are involved, our finding that *Atrx* deletion increased soft tissue sarcoma susceptibility to dsDNA oncolytic herpesvirus *in vitro* suggests that patients with sarcomas and perhaps other tumors with *ATRX* loss-of-function mutations may benefit from oncolytic virus (OV) therapy. OV therapy is an emerging cancer therapy, utilized in more than 90 recent clinical trials (47). Most of the trials underway are utilizing either oncolytic adenovirus, herpesvirus, polio virus, or vaccinia virus (48–50). In fact talimogene laherparepvec (T-VEC), an oncolytic herpesvirus similar to the oncolytic herpesvirus used in this study, was the first OV therapy approved by the FDA to treat unresectable metastatic melanoma (29, 50, 51). Because deletion of *Atrx* sensitizes sarcoma cells to oncolytic herpes virus and radiation therapy, tumors with loss-of-function mutations in *ATRX* may be particularly sensitive to the combination of oncolytic herpesvirus with radiation therapy. In addition, a recently completed phase 2 clinical evaluating a PD-1 immune checkpoint inhibitor and T-VEC oncolytic herpesvirus combination therapy in advanced sarcoma showed a promising overall objective response rate of 35% (52). Our work lays the foundation for future studies examining the impact of ATRX loss on the response of soft tissue sarcomas to therapeutic combinations of PD-1 inhibitors, oHSV, and radiation.

Collectively, these results show that loss of ATRX function impairs the cGAS-STING signaling pathway and promotes the response to radiation therapy and oncolytic virus therapy in soft tissue sarcoma. These findings suggest that for these cancer therapies *ATRX* mutation status in sarcomas and perhaps other cancers may be a biomarker for treatment response.

## Supporting information

Supplemental Table 1: Key Resources

Supplemental Table 2: Enrichment Pathways

## Acknowledgements

We thank Andrea Daniel for her assistance in experimental design and for providing feedback on the manuscript. We thank Richard Gibbons for providing the ATRX^FL^ mice, Tyler Jacks for the LSL-Kras^G12D^ mice, Anton Berns for the p53^FL^ mice, and Chen-Ming Fan for the Pax7-CreER mice. Graphics for schematics were created using Biorender.com. This work was supported by the National Cancer Institute of the US, NIH (award nos. F30CA232652 and 3R35CA197616-04S1 to W.F. and R35CA197616 to D. G. K.)

## Authors’ Contribution

Conceptualization: W.F. and D.G.K. Methodology. W.F., S.B.D., J.S., W.E., and D.G.K. Formal Analysis: W.F., D.L.C., and D.M.C. Investigation: W.F., M.P., C.S., L.L., K.D., D.Z., Y.M., and N.T.W. Resources: A.J.W., S.B.D., J.S. and W.E. Writing-Original Draft: W.F. and D.G.K. Writing Review and Editing: W.F., K.D., A.J.W., D.M.C. and D.G.K. Visualization: W.F. Supervision: D.G.K

---

**Supplementary Table 1: Reagents**

Detailed list of antibodies, primers, chemotherapeutic agents, and other materials used in this work.

**Supplementary Table 2: Significant Pathway Enrichments**

Statistically significant pathway enrichment results for samples described in this paper.

## Methods

### Data Code and Availability

- RNA-sequencing data has been submitted to Gene Expression Omnibus at NCBI, and is publicly available under GEO accession number GSE167537.

### Experimental Models and Subject Details

#### Mouse Strains

Mouse strains used in the primary mouse model were on a mixed 129S_4_/SvJae and C57/Bl6 background. *LSL-Kras^G12D^* mice were provided by Tyler Jacks (Massachusetts Institute of Technology) (53). *p53^Fl^* mice were provided by Anton Berns (Netherlands Cancer Institute, Amsterdam, Netherlands) (54). *Atrx^Fl^* mice were provided by Richard Gibbons (University of Oxford) (55). Pax7^CreER^ were provided by Cheng-Min Fan (56). NCre nude mice were purchased from Taconic Biosciences. To minimize the effects of genetic background, littermate controls were used in experiments to ensure any genetic modifying variables would be randomly distributed between experimental and control groups. Mice were randomly assigned into experimental treatment cohorts within cages to ensure balancing of sex and age. All mice used in experimental cohorts did not receive any treatment or undergo any procedures prior to start of the experiments described in this paper.

#### Generation of Isogenic Cell Lines and Cell Culture

pSP Cas9(BB)-2A-GFP (PX458) with ATRX gRNA or vector control were transfected into newly derived primary soft tissue sarcoma cell lines (KP) using the TransIT-LT1 transfection reagent (MIR2304, Mirus). Forty-eight hours after transfection, EGFP positive cells were single cell sorted via fluorescence activated cell sorting into 96 well plates using either an Astrios Cell Sorter (Beckman Coulter) or a BD Diva sorter (BD). For vector control, PX458 plasmid containing GFP and Cas9 but without insertion of a gRNA was transfected and identical GFP screening was performed. Cell lines from each single cell clone were then screened for loss of *ATRX* protein expression by western blot. Antibodies used for western blot and immunofluorescence are listed in Supplemental Table 1. ATRX WT control cell lines underwent all screening steps concurrently with their ATRX KO isogenic cell line counterparts. Passaging of sarcoma cell lines was performed in DMEM (Thermo Fisher Scientific, 11965092) supplemented with 10% fetal bovine serum (Thermo Fisher Scientific, 16000044), 1% Pen-Strep (Thermo Fisher Scientific, 11965092), and 1% L-glutamine (Thermo Fisher Scientific, A2916801). Cell lines were incubated at 37°C with 5% CO_2_ in a humidified cell culture incubator.

#### Sarcoma Induction

Primary sarcomas were generated in Pax7^CreER^; *LSL-Kras^G12D^*; *p53^Fl/Fl^* (P7KP) and *Pax7^CreER^*;*LSL-Kras^G12D^*; *p53^Fl/Fl^*; *Atrx^Fl^* (P7 KPA) mice as previously described (57). To activate CreER in a spatially and temporally controlled manner in Pax7-expressing muscle satellite cells, 4-hydroxytamoxifen (4-OHT, Sigma, H7904) was dissolved in 100% DMSO at a concentration of 20mg/ml and 25 ul of the 4-OHT solution was injected via intramuscular injection into the left mouse gastrocnemius muscle. For P7 P + MCA and P7 PA + MCA models, 4-OHT intramuscular injection was followed within 30 minutes by injection into the same muscle with of 300 µg MCA (Sigma) resuspended in sesame oil (Sigma) at 6 µg/µL. Sarcoma cell lines for *in vitro* deletion of *Atrx* by CRISPR/Cas9 were generated in *LSL-Kras^G12D^*; *p53^Fl/Fl^* (KP) mice on a 129 Sv/Jae or mixed 129 Sv/Jae and C57/Bl6 background by injection of Cre-expressing adenovirus into the gastrocnemius muscle as previously described (22). KP *Atrx* WT and KP *Atrx* KO transplant sarcomas were generated by injecting 50,000 cells in a 1:1 mixture of DPBS (Gibco) and high concentration matrigel (Corning) into the gastrocnemius muscle. Mice were anaesthetized with 2% isoflurane prior to all injections or procedures.

#### Radiation Therapy

When tumors reached 50-250 mm^3^ (Day 0, D0) mice were randomized to treatment groups, then tumor growth was monitored via three times weekly caliper measurements in two dimensions. Sarcoma irradiation was performed using the Precision Xrad 225 Cx small animal image-guided irradiator (58). The radiation therapy field was centered on the mouse gastrocnemius tumor via fluoroscopy with a 40 kilovolt peak (kVp), 2.5 mA x-rays using a 0.3 mm copper filter. Sarcomas were irradiated with parallel-opposed anterior and posterior fields with an average dose rate of 300 cGy/min prescribed to a midplane with 225 kVp, 13 mA x-rays using a 0.3-mm copper filter and a collimator with a 40×40 mm^2^ radiation field at treatment isocenter. Mice were euthanized using CO2 if they met their humane endpoint as determined by the IACUC protocol (moribund or in distress), or when tumor volume exceeded 2cm^3^, in accordance with Duke IACUC guidelines. Mice that were sacrificed for non-tumor related reasons were excluded from analysis. Graphpad Prism version nine was used for non-linear fit modeling of tumor growth after irradiation. For this modeling the equation (Y=Y_0_*exp(k*X) was used, Y_0_ was constrained to be shared between the two datasets and to fall between the 50-250 mm^3^. Convergence criteria was set as “strict” and 10,000 interations were run using the least squares regression fitting method.

#### Immunohistochemistry

Tumors were fixed in 10% neutral buffered formalin overnight, transferred to 70% ethanol, paraffin-embedded, and sectioned to 5 *μ*m thickness. Deparaffinized and rehydrated slides were blocked with 3% hydrogen peroxide. Antigen retrieval was performed by boiling in 2% citrate-based antigen unmasking solution (VECTOR H-3300) for 15 min. Slides were blocked with 5% normal serum in PBS + 0.25% Tween-20 and incubated overnight at 4C with primary antibodies as listed in Supplementary Table 1. Following this step, slides were diluted in PBS + 0.25% Tween-20 + 5% normal serum overnight at 4°C. After washing, slides were incubated with biotinylated secondary antibodies, including 1:200 horse anti-mouse IgG (VECTOR BA-2000) or 1 : 200 goat anti-rabbit IgG (VECTOR BA-1000) for 1 hour at room temperature. Slides were incubated with VECTASTAIN Elite ABC Reagent (VECTOR PK-7100) for 30 min at room temperature before signal was visualized with the 3,3′-diaminobenzidine (DAB) peroxidase substrate kit (VECTOR SK-4100). Representative images were acquired with a Leica DFC450 brightfield microscope using Leica Suite software (Leica Microsystems).

Hematoxylin and eosin (H&E) stained tissue sections were examined by a sarcoma pathologist (D.M.C.) blinded to genotype and treatment. For quantification of tumor necrosis, the sarcoma pathologist outlined areas of necrosis. ImageJ was used to quantify percentage of tissue that was identified as necrotic. Tumors type was classified by D.M.C. based on the full panel of sarcoma classification markers that utilized 9 antibodies (Supplementary Table 1). For Ki-67 IHC experiment, ImageJ was used to quantify positively stained cells by a single observer blinded to genotype and treatment.

#### In vitro Immunocytochemistry

Nunc Lab-Tek II Chambered Coverglass was used to grow and image cells (ThermoFischer Scientific, 155379). Cells were fixed with 4% paraformaldehyde, blocked and washed using the MAXpack Immunostaining Media Kit (Active Motif), then incubated with primary antibody as listed in Supplementary Table 1 overnight at 4°C. Secondary antibody (Supplementary Table 1) was incubated for 1 hour at room temperature. Cells were stained with DAPI and then Prolong Diamond Antifade Mountant. Imaging was performed on a Leica SP5 Inverted Confocal Microscope, with consistent settings to allow for comparison. For each experiment a minimum of 50 cells were imaged for each experiment, then analyzed by a researcher blinded to treatment and genotype using Leica Suite software (Leica Microsystems). Data from these experiments represent the average of at least three independent experimental technical replicates.

#### TUNEL Assay In Situ Cell Death Detection Kit, AP (Roche, #11684809910)

Frozen Tissue sections were allowed to thaw at room temperature for 5 minutes, then sections were fixed in cold 4% paraformaldehyde for 10 minutes. Samples were washed twice with ice cold PBS, then incubated with 0.1% Triton X-100 and 0.1% sodium citrate in water (made fresh) for five minutes. Samples were washed 3 times for 5 minutes each in PBS. Then 100 microliter TUNEL reaction mixture was added to each sample and incubated for 1 hour at 37°C. Each sample was washed three times with PBS, mounted, and then imaged with a DFC340FX fluorescence microscope using Leica Software.

#### Mitotic Dysfunction Assays

Micronuclei were defined as regions of DAPI staining located in close proximity to a nucleus with a size between approximately 1/3 and 1/64 the size of a nucleus, which could be determined to be distinct from the nucleus. Chromosome bridges were defined as occurrences of a single thin clear line connecting two otherwise separate nuclei. TIF co-localization was defined by visual inspection in multiple channels and were counted if there was an instance of a 53BP1 focus that was contiguous or overlapping with a telomere focus. Only a single TIF event was called per 53BP1 focus, even if multiple telomere foci were present. Mitotic catastrophe was identified by the presence of multiple micronuclei that were associated with a single nucleus (59–61).

#### ImmunoFISH

Telomere ImmunoFISH was performed as described in (62) (Basic Protocol 1), with the following modifications: Nunc Lab-Tek II Chambered Coverglass was used to grow and image cells (ThermoFischer Scientific, 155379). Cells were incubated for 30 seconds in pre-extraction buffer, and Alexa-Fluor 488 conjugated telG FISH probe was used (PNA BIO). For the FISH hybridization step, slides were preheated for 1 minute before adding the telomere probe to minimize background staining. Imaging was performed on a Leica SP5 Inverted Confocal Microscope, and imaging for each experiment was performed using consistent settings to allow for comparison.

#### Cell Line Generation from Primary Sarcomas

Tumors were collected immediately after mouse sacrifice. Tumor tissue was minced and digested in dissociation buffer in PBS (Thermo Fisher Scientific, 14040133) containing collagenase type IV (5 mg/ml, Thermo Fisher Scientific, 17104-019), dispase (1.3 mg/ml, Thermo Fisher Scientific, 17105-041) as well as trypsin (0.05%, Thermo Fisher Scientific, 25200056) for about 1 h at 37°C in a cell culture hood and incubator. Cells were washed with PBS (Thermo Fisher Scientific, 10010023) and filtered using a 40 *μ*m sieve (Corning, 431750) and cultured for at least five passages to deplete stroma before being used for experiments. To assess recombination of the *Atrx* floxed allele, tumor cell line genomic DNA was isolated and genotyped by PCR to amplify exon 18 or the recombined *Atrx^FL^* allele using primer sets in Supplementary Table 1.

#### IC50 assays

For IC50 assays, cells were plated at 1000 cells per well in a 96 well plate, then treated with 1:2 serial dilutions of doxorubicin. Cells were incubated in a cell culture incubator with 5% carbon dioxide at 37 degrees Celcius for 72 hours before cells were washed twice with PBS then measured via CellTiter-Glo (Promega). IC50 experiments were repeated three times for each cell line shown.

#### Genotyping

Mice were genotyped using tails collected from mouse pups. Tail genomic DNA was extracted using a KAPA mouse genotyping Kit (KAPA Biosystems KK7352), and PCR was performed using primers as described in Supplementary Table 1. Additional genotyping was performed by shipping mouse tails to Transnetyx, where DNA was extracted and analyzed using a Taqman qpcr assay. For *Atrx* exon 18 assay (Figure 2E, top), presence of un-recombined exon 18 is shown by the presence of a band at 1000 bp, while recombination is visualized by no band being present. For *Atrx* recombination assay, *Atrx* WT can be visualized as a 1400 bp band, floxed *Atrx* as a 1600 bp band, and recombined *Atrx* as a 600bp band.

#### Immunoblotting

Samples were lysed in RIPA buffer for 30 min on ice (Sigma-Aldrich, R0278), sonicated briefly, and then centrifuged at 10,000x·g for 10 min at 4°C. For western blots analyzing ATRX, lysate was not frozen, as this resulted in a marked reduction of signal. Protein concentration was determined for the lysate supernatant by BCA assay (Pierce, 23225). Samples were boiled in 4x Laemmli sample buffer (Bio-Rad, 1610747) before loading in a 4–20% Tris-glycine polyacrylamide gel. Membranes were blocked in 1:1 TBS:Intercept blocking buffer (LiCor) for 1 hour. Next, the membranes were incubated overnight at 4°C with primary antibodies diluted in TBS-T (0.1% Tween-20) at concentrations as indicated in Supplementary Table 1. The membranes were washed three times in TBS-T before secondary antibody incubation with goat anti-rabbit IRDye800 (Li-Cor Biosciences, P/N 925-32211) and goat anti-mouse IRDye680 (Li-Cor Biosciences, P/N 925-68070) both at 1:10,000 dilutions in TBS-T for 1 h at room temperature. The membranes were washed three times in TBS-T and imaged using an Odyssey CLx (Li-Cor Biosciences). Image analysis for normalization and quantification was performed using Image Studio (version 5.2, Li-Cor Biosciences, P/N 9140-500).

#### RT-PCR

Cells were lysed with TRIzol reagent (Thermo Fisher, 15596026). RNA was isolated from samples using a Direct-zol RNA MiniPrep kit (Zymo Research, R2051). RNA samples were reverse transcribed to cDNA using an iScript Advanced cDNA Synthesis Kit (Bio-Rad, 1725038). TaqMan probes from Thermo Fisher were used for RT-PCR: *IfnB1* (Mm00439552), *Gapdh* (Mm99999915), and *Tbp* (Mm01277042_m1). Taqman fast advanced master mix was used for the qPCR assay (ThermoFischer Scientific 4444556). Plates were run on a QuantStudio 6 Flex Real-Time PCR System (Thermo Fisher), and data were analyzed using the comparative C_T_ method to generate expression fold-change values. *Gapdh* or *Tbp* expression was used as an internal control for RNA concentration across samples. Every sample was run in triplicate and results were averaged for each assay.

#### RNA Sequencing Sample Generation and Barcoding

RNA sequencing was performed by Novogene. RNA degradation and contamination was monitored on 1% agarose gels, and RNA purity was checked using the NanoPhotometer spectrophotometer (IMPLEN, CA, USA). RNA integrity and quantitation were assessed using the RNA Nano 6000 Assay Kit of the Bioanalyzer 2100 system (Agilent Technologies, CA, USA). A total amount of 1 μg RNA per sample was used as input material for the RNA sample preparations. Sequencing libraries were generated using NEBNext® UltraTM RNALibrary Prep Kit for Illumina® (NEB, USA) following manufacturer’s recommendations and index codes were added to attribute sequences to each sample. mRNA was purified from total RNA using poly-T oligo-attached magnetic beads. Fragmentation was carried out using divalent cations under elevated temperature in NEBNext First Strand Synthesis Reaction Buffer (5X). First strand cDNA was synthesized using random hexamer primer and M-MuLV Reverse Transcriptase (RNase H-). Second strand cDNA synthesis was subsequently performed using DNA Polymerase I and RNase H. Remaining overhangs were converted into blunt ends via exonuclease/polymerase activities. After adenylation of 3’ ends of DNA fragments, NEBNext Adaptor with hairpin loop structure were ligated to prepare for hybridization. In order to select cDNA fragments of preferentially 150∼200 bp in length, the library fragments were purified with AMPure XP system (Beckman Coulter, Beverly, USA). Then 3 μl USER Enzyme (NEB, USA) was used with size-selected, adaptor ligated cDNA at 37 °C for 15 min followed by 5 min at 95 °C before PCR. Then PCR was performed with Phusion High-Fidelity DNA polymerase, Universal PCR primers and Index (X) Primer (NEB). PCR products were then purified (AMPure XP system) and library quality was assessed on the Agilent Bioanalyzer 2100 system.The clustering of the index-coded samples was performed on an Illumina Novaseq. After cluster generation, the libraries were sequenced on the same machine and paired-end reads were generated.

#### Reads mapping to the reference genome

Reference genome and gene model annotation files were downloaded from genome website browser directly. Indexes of the reference genome were built using STAR and paired-end clean reads were aligned to the reference genome using STAR (v2.5). STAR used the method of Maximal Mappable Prefix (MMP) which can generate a precise mapping result for junction reads. Gene expression counts were generated using the HTSeq algorithm.

#### Bioinformatic Analysis

Raw counts from Novogene’s processing pipeline were obtained from the company. For cell line RNA seq analysis, genes were first filtered if they did not have at least 10 reads in a single sample from each of the 3 replicates. Normalization and differential expression were carried out using the DESeq2 Bioconductor package with the R statistical programming environment. In each analysis, ‘replicate’ was used as a cofactor in the model. The false discovery rate was calculated to control for multiple hypothesis testing. Gene set enrichment analysis (63) was performed to identify gene ontology terms and pathways associated with altered gene expression for each of the comparisons performed

#### Immune Infiltrate Estimation

For each tumor sample, gene TPM were collected into a spreadsheet and then input into http://timer.cistrome.org/ for immune estimation (64, 65). Data output from the mMCP Counter algorithim for T-cell infiltrate score and Macrophage/Monocyte infiltrate score were then collected for the samples and compared using Graphpad prism (26).

#### Cancer genomic database analysis

For determination of the frequency of ATRX mutation in human cancers, the TCGA database was queried via cBioportal. For overall prevalence of ATRX in human cancer, “Curated set of non-redundant studies” was selected, and was queried for genomic profile of mutations and copy number alterations in ATRX. For data on frequency and position of ATRX mutations in soft tissue sarcoma, “Sarcoma (TCGA, PanCancer Atlas)” study was selected, UPS, LMS and MFS sarcoma subtypes were selected, then were queried for genomic profile of mutations and copy number alterations in ATRX. Disease free survival data was obtained via CBioportal. Whole genome sequencing mutation and phenotype data of undifferentiated pleomorphic sarcomas was accessed from (2). Samples harboring frameshift or missense mutations in ATRX exons, as well as samples with copy number deletion, fusions or inversions were classified as “ATRX altered”.

#### Statistics

For bar graphs, all data are presented as means +/− SEM. Student’s t-test (two tailed with Welch’s correction) was used to compare the mean of two groups unless analyzing results for multiple paired isogenic cell lines. For statistical analysis of multiple paired isogenic cell lines, a ratio paired t-test was performed. Graphs with multiple comparisons were analyzed using a two-way ANOVA with Tukey’s multiple comparisons test. For tumor growth experiments survival curves were estimated for each group, considered separately, using the Kaplan-Meier method and compared statistically using the log-rank test. A p value less than 0.05 indicated significance. Prism 9 (GraphPad Software, Inc.) was used for statistical analysis.

### Study Approval

All animal studies were approved by the institutional Animal Care and Use Committee (IACUC) at Duke University.

---

**Supplementary Figure 1:**
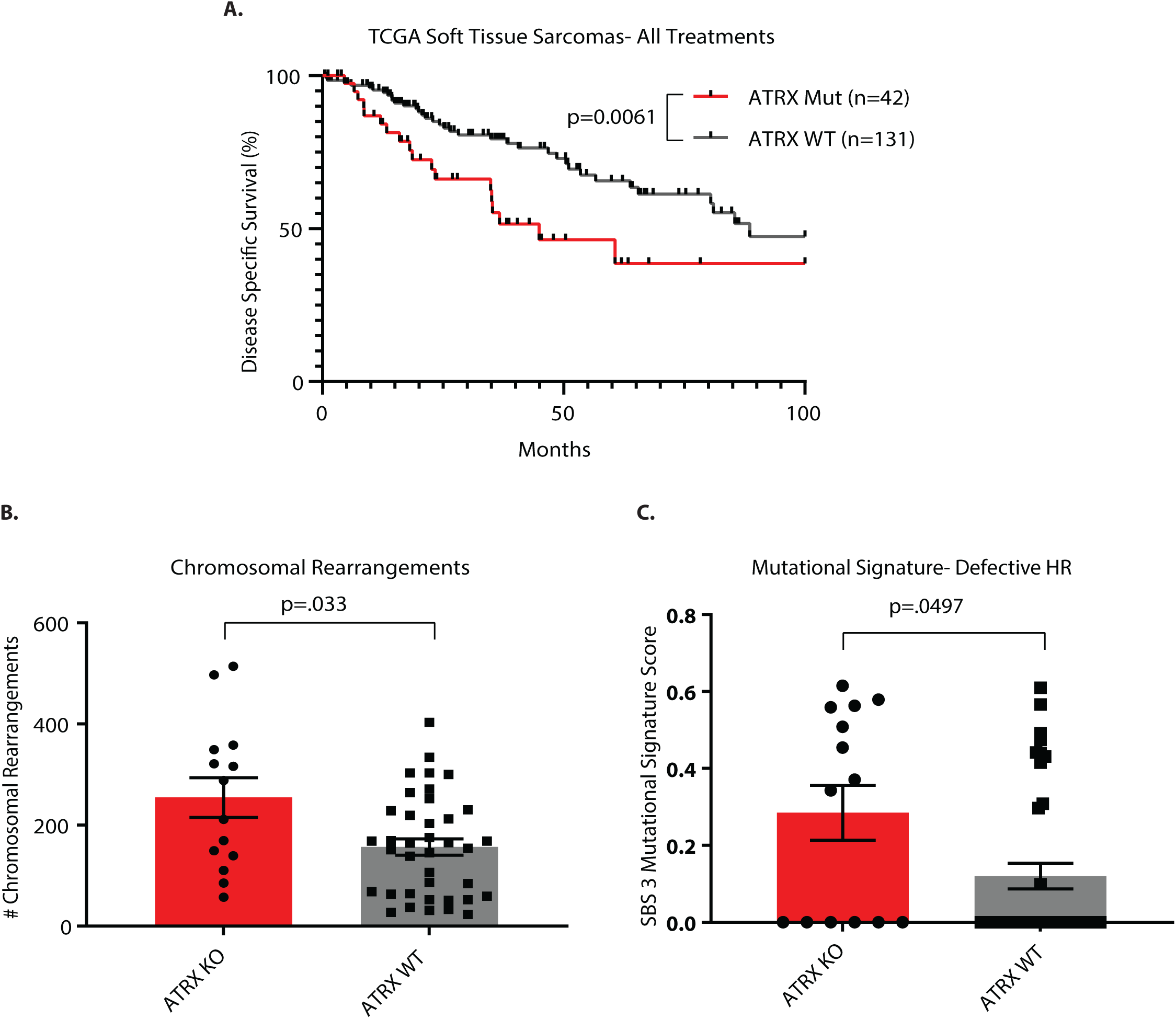
Characteristics of Human Soft Tissue Sarcomas. A) Disease specific survival for a cohort of soft tissue sarcomas (leiomyosarcomas, myxofibrosarcomas, and undifferentiated pleomorphic sarcomas), comparing sarcomas with ATRX genomic alteration (ATRX Mut, n=42) to sarcomas without ATRX genomic alteration (ATRX WT n=131). Statistical comparison was performed using a log-rank (Mantel Cox) test. B) Analysis of whole genome sequencing of undifferentiated pleomorphic sarcoma showing average number of chromosomal rearrangements in 14 *ATRX* altered and 38 *ATRX* wild type (WT) human tumors. C) Analysis of whole genome sequencing of undifferentiated pleomorphic sarcoma showing COSMIC mutation signature 3 score, which reflects defective homologous recombination (HR)-based double strand DNA break repair, in 14 ATRX altered and 38 ATRX WT human tumors. Statistical analysis for B) and C) was performed using a Welch’s t-test. Each data point represents a biological replicate.

**Supplementary Figure 2:**
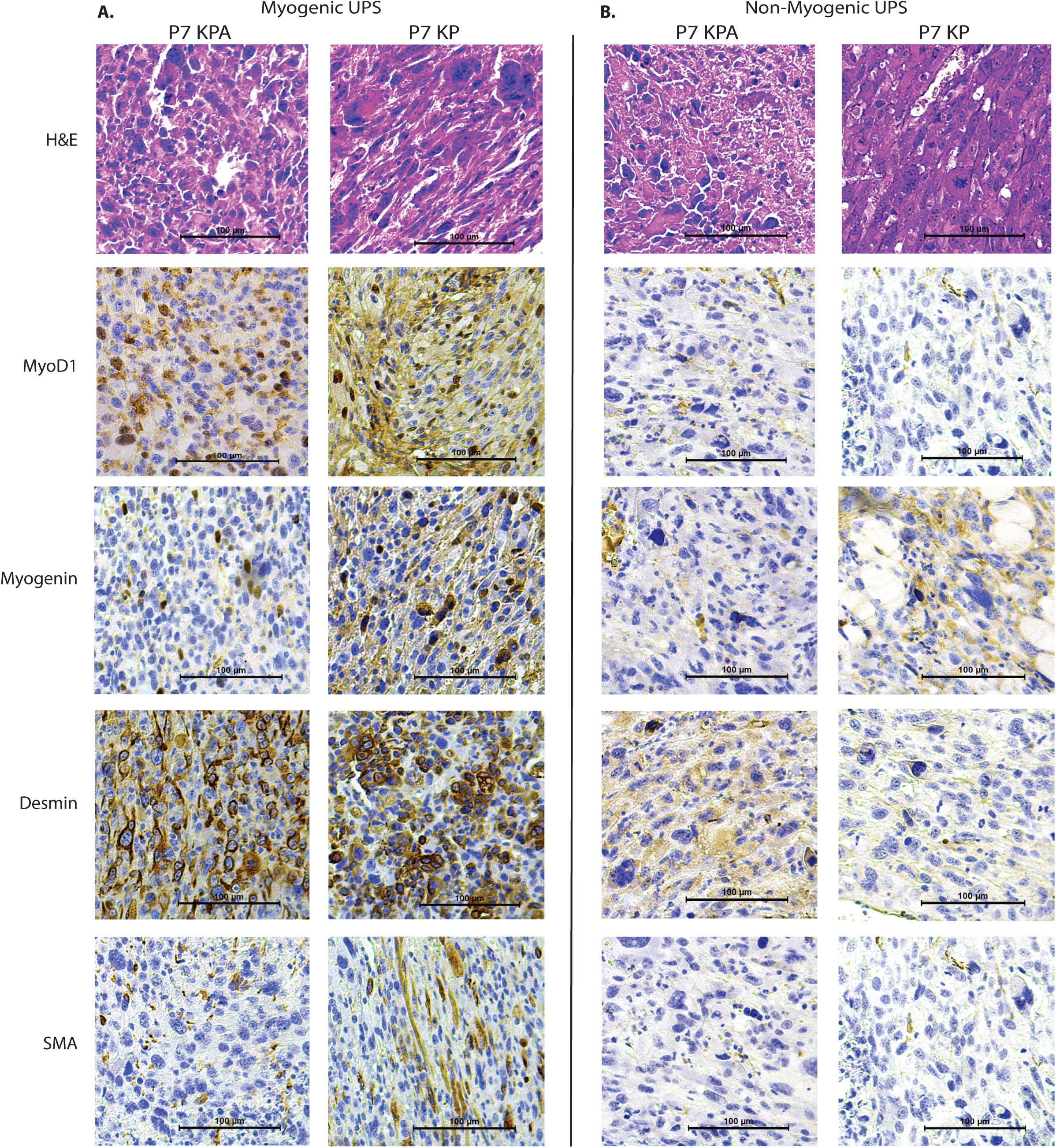
Immunohistochemistry for classification of soft tissue sarcomas. A-B) Representative H&E staining and immunohistochemistry for myogenic markers for formalin fixed, paraffin embedded tumors used to classify soft tissue sarcomas from the P7 KPA (left) and P7 KP (right) mice. Myogenic UPS H&E staining images are identical to images shown in figure 2. H&E staining as well as immunohistochemistry of muscle markers MyoD (nuclear), Myogenin (nuclear), Desmin (cytoplasmic) and SMA (cytoplasmic) for myogenic UPS (A) and non-myogenic UPS (B). Tumors shown were classified as based on the full panel of sarcoma classification markers that utilized 9 antibodies (Supplementary Table 1). Scale bar is 100 μm.

**Supplementary Figure 3:**
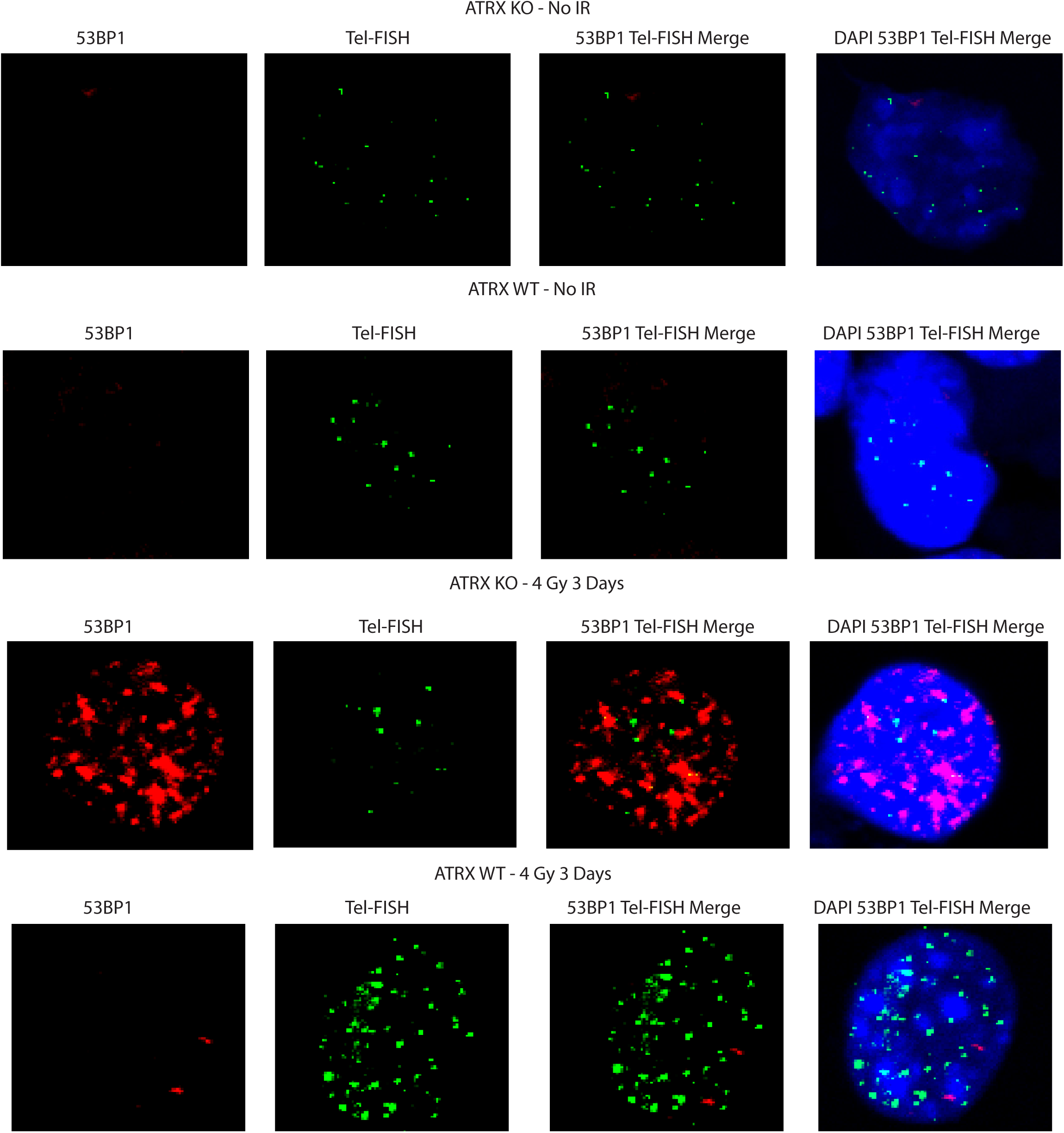
ImmunoFISH staining. Representative images showing immunofish staining of an ATRX KO and WT isogenic cell line pair. Cells were fixed and stained 3 days after 4 Gy irradiation or no irradiation. Each series shows 53BP1 staining (red), telomere FISH staining (green), merged image of 53BP1 and telomere fish staining, and merge of 53BP1, telomere FISH, and DAPI staining.

**Supplementary Figure 4:**
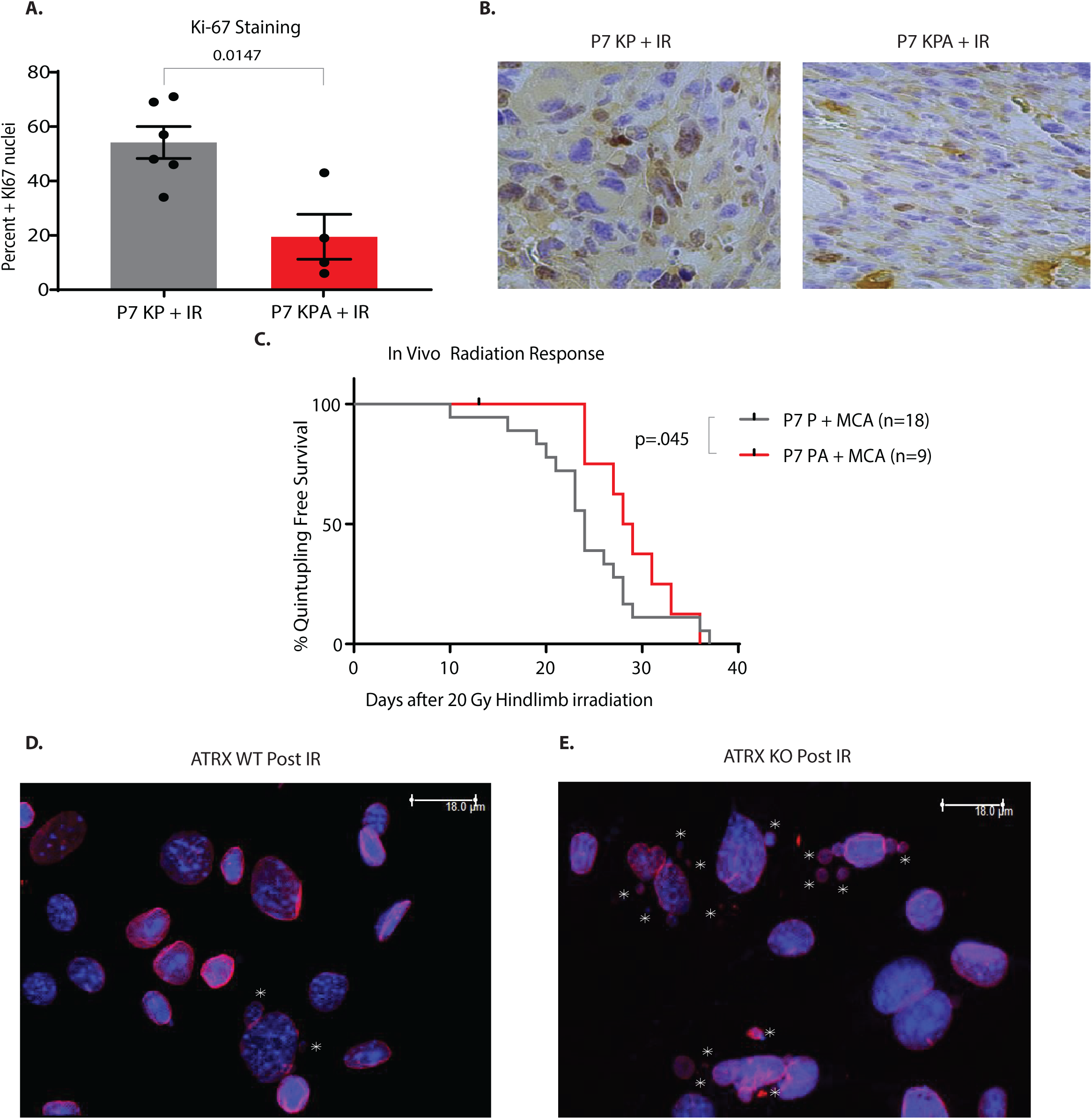
Effect of *Atrx* deletion on radiation response in soft tissue sarcomas. A) Quantification of Ki67 staining from sarcomas from P7 KP and P7 KPA mice harvested 6 Days after 20 Gy. Each data point represents a biological replicate. B) Representative Ki67 staining from sarcomas from P7 KP (left) and P7 KPA (right) mice harvested 6 days after 20 Gy. C) Time to tumor quintupling after treatment with 20 Gy in the P7 P + MCA primary mouse model of soft tissue sarcoma. D-E) Representative images of micronuclei in ATRX WT (D) and ATRX KO (E) sarcoma cells three days after 4 Gy as quantified in Figure 6A. DAPI is blue and Lamin B1 is red. The presence of nuclear envelope enclosed micronuclei can be observed (white stars).

**Supplementary Figure 5:**
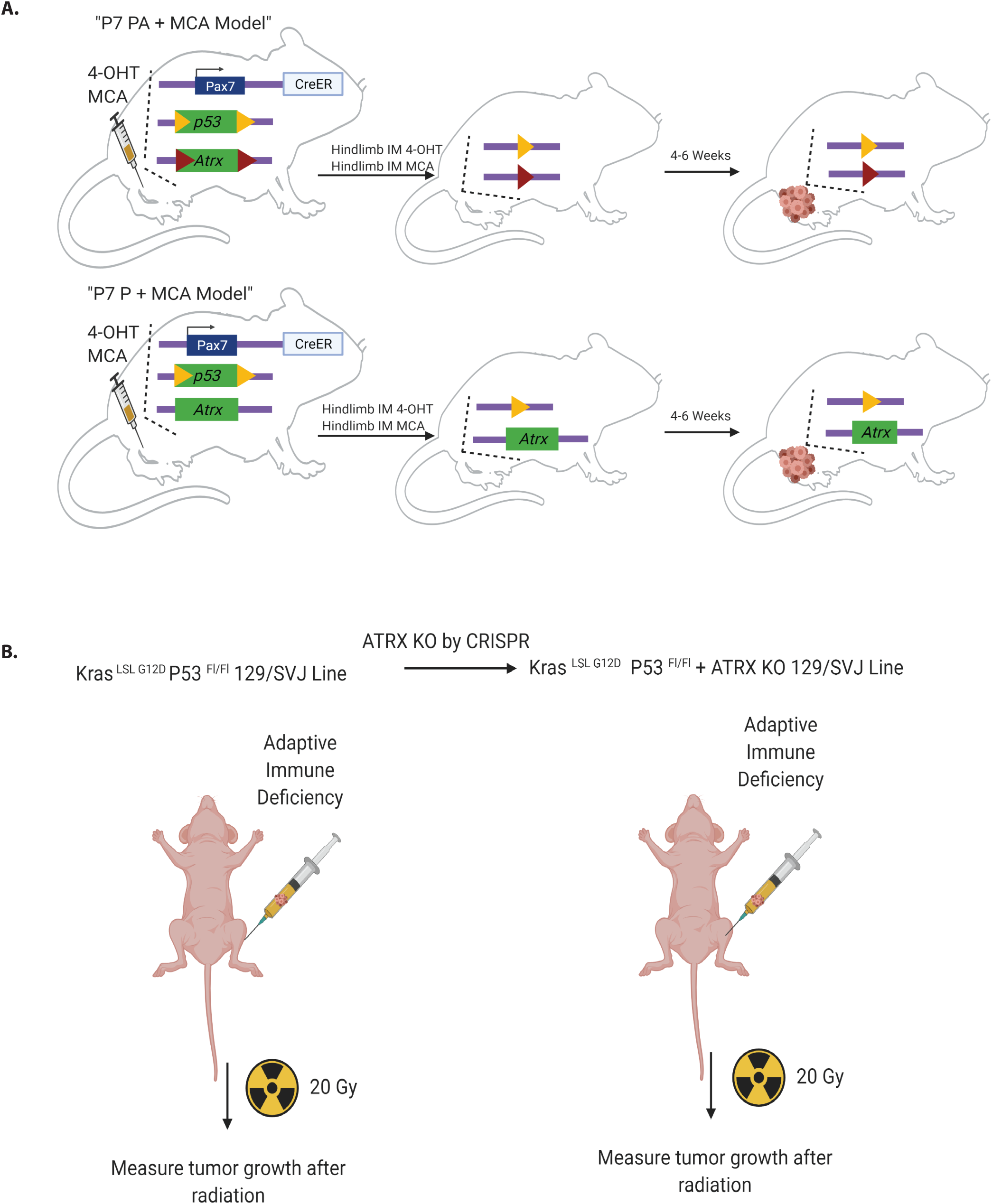
Experimental schematics for mouse experiments. A) Diagram showing the P7 PA model of soft tissue sarcoma, triangles represent loxp sites. Mice receive intramuscular injection of 4-OHT in DMSO followed within 30 minutes by injection of MCA into the same muscle. The P7 P + MCA model does not utilize a genetically engineered conditional oncogenic *Kras^G12D^* allele, but instead is initiated by intramuscular injection of 4-OHT in *Pax7-CreER^T2^*; *p53^FL/FL^* mice to delete both *p53* alleles followed by injection of 3-methylcholanthrene, a potent carcinogen that drives base substitutions. The P7 PA + MCA model is identical except for the presence of loxp sites flanking exon 18 of *Atrx* that allows deletion of both ATRX and *p53* following administration of 4-OHT. B) Diagram showing the design of tumor transplant experiment into immunodeficient mice. Nude mice received hindlimb intramuscular injection of the gastrocnemius with 50,000 cells in matrigel. After tumors were detected by caliper, mice were treated with 20 Gy and measured three times a week until tumors reached 2000 mm^3^ or was required by IACUC humane endpoints.

**Supplementary Figure 6:**
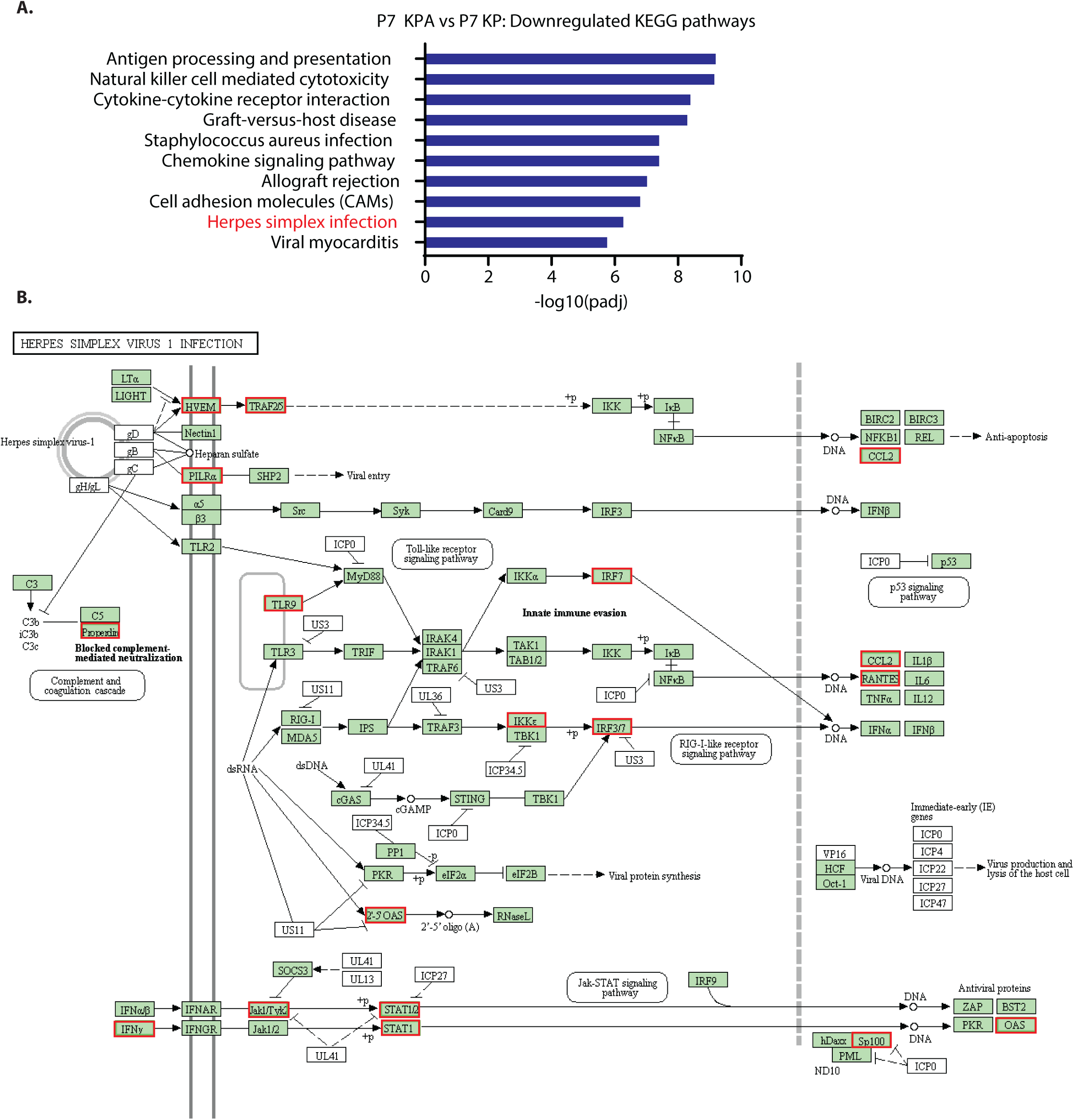
ATRX deletion results in reduced expression of HSV-1 innate immune defense pathways. A) Top ten results for KEGG enrichment analysis of RNA sequencing comparing unirradiated P7KPA sarcomas (n=4) to unirradiated P7 KP tumors (n=4). B) KEGG pathway map for cellular response to HSV-1 infection. Cellular genes are in green, HSV genes are in white. Red boxes are used to indicate genes that are downregulated (P7KPA No IR vs P7KP No IR) in the KEGG pathway, no genes were upregulated in this KEGG pathway.

